# Perfusable biohybrid designs for bioprinted skeletal muscle tissue

**DOI:** 10.1101/2023.01.12.523732

**Authors:** Miriam Filippi, Oncay Yasa, Jan Giachino, Reto Graf, Aiste Balciunaite, Robert K. Katzschmann

## Abstract

Engineered, centimeter-scale skeletal muscle tissue (SMT) can mimic muscle pathophysiology to study development, disease, regeneration, drug response, and motion. Macroscale SMT requires perfusable channels to guarantee cell survival and support elements to enable mechanical cell stimulation and uniaxial myofiber formation. Here, stable biohybrid designs of centimeter-scale SMT are realized via extrusion-based bioprinting of an optimized polymeric blend based on gelatin methacryloyl and sodium alginate, which can be accurately co-printed with other inks. A perfusable microchannel network is designed to functionally integrate with perfusable anchors for insertion into a maturation culture template. The results demonstrate that (*i*) co-printed synthetic structures display highly coherent interfaces with the living tissue; (*ii*) perfusable designs preserve cells from hypoxia all over the scaffold volume; and (*iii*) constructs can undergo passive mechanical tension during matrix remodeling. Extrusion-based multimaterial bioprinting with our inks and design realizes *in vitro* matured biohybrid SMT for biomedical, nutritional, and robotic applications.

## 1. Introduction

Skeletal muscle tissue (SMT) is the largest organ by mass in the human body, and it is crucial for locomotion, posture, respiration, physiology, and energy homeostasis.^1,2^ Genetic or acquired diseases, as well as damage and aging can render this organ dysfunctional and cause significant impairment in human quality of life and health.^4^ In the last three decades, several three-dimensional (3D) SMT culture systems have been developed to mimic the native muscle microenvironment and understand skeletal muscle function, plasticity, and disease.^1,5^ This understanding not only improved cell culture models for biomedical research^6,7^ and engineered tissue grafts for medical implantation,^2,8,9^ but also enabled controllable biomachines with dynamic abilities (*i*.*e*., biohybrid robots).^10–12^

Through the years, SMT was engineered *in vitro* via approaches that aim at replicating the ultrastructure of native muscle tissue, which is composed of highly co-oriented myofibers. Topographical, chemical, and electrical cues can align adjacent myoblasts to generate uniaxially oriented myotubes.^13–20^ These local cues deliver stimuli to cells when these cells grow on surfaces, in planar geometries (*e*.*g*., muscular thin films), or in microtissues.^21–24^ However, they can not be easily applied to block-like 3D SMT constructs over the centimeter scale, as the stimuli would hardly reach the cells residing in the internal core of the constructs, leading to inhomogeneous cell stimulation.

In contrast to local cues, applying mechanical stress to a scaffold filled with myoblasts is an effective way to form myotubes and control the orientation of the forming fibers all over a scaffold’s volume.^3^ To convey stress and strain to cells, one can use biohybrid tissue configurations that interface the biological tissue with synthetic elements. Biohybrid SMT constructs contain synthetic structures, such as tendon-mimicking anchors and skeletons, to counteract forces occurring during muscle tissue maturation. These forces arise from the shrinkage of the hydrogels that are used as cell scaffolding materials and the remodeling action of cells on the scaffolding materials. Biohybrid designs simplify the mechanical stimulation of muscle progenitor cells to form intact co-oriented assemblies of fibers in cm-scaled 3D models.

Despite their utility in engineering millimeter-scale SMT,^3,10,25^ biohybrid designs have not yet been adapted for the optimal development of SMT constructs at the centimeter scale. The two essential challenges in scaling up SMT designs are related to (*i*) the perfusion and (*ii*) the structural stability of the final constructs. First, biohybrid designs should allow for the efficient distribution of nutrients and oxygen to guarantee cell survival. Second, biohybrid designs should feature coherent interfaces between the muscle tissue and the synthetic elements; these interfaces increase the stability of constructs during *in vitro* culture and reduce force dispersion that would otherwise occur during the muscle contraction if there is a mismatch in adherence between different materials. A suitable design and fabrication strategy are needed to enable perfusion and structural stability within a construct.

Bioprinting technologies can position biomaterials, cells, and bioactive factors in a single construct to achieve architectures that mimic native tissues.^26^ SMT constructs (mm^3^–cm^3^ scale) have been fabricated via extrusion-based 3D bioprinting and used as pathophysiological models, implantable grafts, and drug screening platforms.^26,27^ For example, Kim et al. co-printed a cell-laden bioink and a sacrificial gelatin-based ink to generate an SMT construct of one cm^3^ in size with channels (∼400-500 μm in diameter) throughout and two supporting anchors at each end.^28^ Tissue maturation and longevity of their SMT were hindered because the anchor’s design obstructed the fluid exchange between the channels and the environment, and no mechanical stressing was applied to the anchors.

Here, we formulated bioinks and applied them to extrusion-based bioprinting to fabricate our biohybrid SMT design that: (*i*) models the functional perfusion of native muscle structure with a high-resolution distribution of vessels, which preserves cell viability within the cm-scale tissue; (*ii*) supports crucial mechanical tensioning for uniaxial fiber growth; and (*iii*) allows for controlled fiber formation and maturation. To realize this maturable muscle biohybrid structure, we optimized the 3D construct design for a stable, yet perfusable, interface of tissue with anchoring structures. Then, we formulated suitable synthetic inks and a bioink, and characterized them for their rheological properties and printability to select the most suitable components for the realization of the biohybrid. We achieved (*i*) perfusable longitudinal microchannels of 200 μm diameters that were finely distributed in the tissue and supported not only cell survival but also alignment homogeneously all over the cm-scale construct, and (*ii*) stable incorporation of synthetic anchors to undergo effective tissue maturation under mechanical tension. Then, we characterized the hypoxic cell responses and biological development of our tissue *in vitro*, and correlated it with data on the perfusion and stability of the biohybrid assembly.

## 2. Results

### 2.1. 3D bioprinting of muscle constructs with structural biomimicry

We used multimaterial extrusion-based 3D bioprinting to realize perfusable, biohybrid SMT (**Figure 1A**). To confer mechanical tension for muscle maturation, the constructs were fixed during the differentiation phase to agar beds at the bottom of culture wells.^29^ Pillars were passed through the anchor holes of the SMT and pierced into the agar (**Figure 1A**). The SMT consisted of a multilayered tissue structure with a nominal size of 3.7×2.4×15 mm and was filled with internal perfusable microchannels with a 200 μm diameter (**Figure 1B**). To generate the channels within the constructs, a sacrificial ink composed of Pluronic F-127 was simultaneously printed with the bioink and then removed after printing by exposing the constructs to a temperature below its gelation point. In addition to the channels, the tissue was co-printed with integrated anchoring structures composed of a mixture of PEGDA and Pluronic F-127, which served to fix the construct to the culture templates used for tissue maturation.

**Figure 1.**
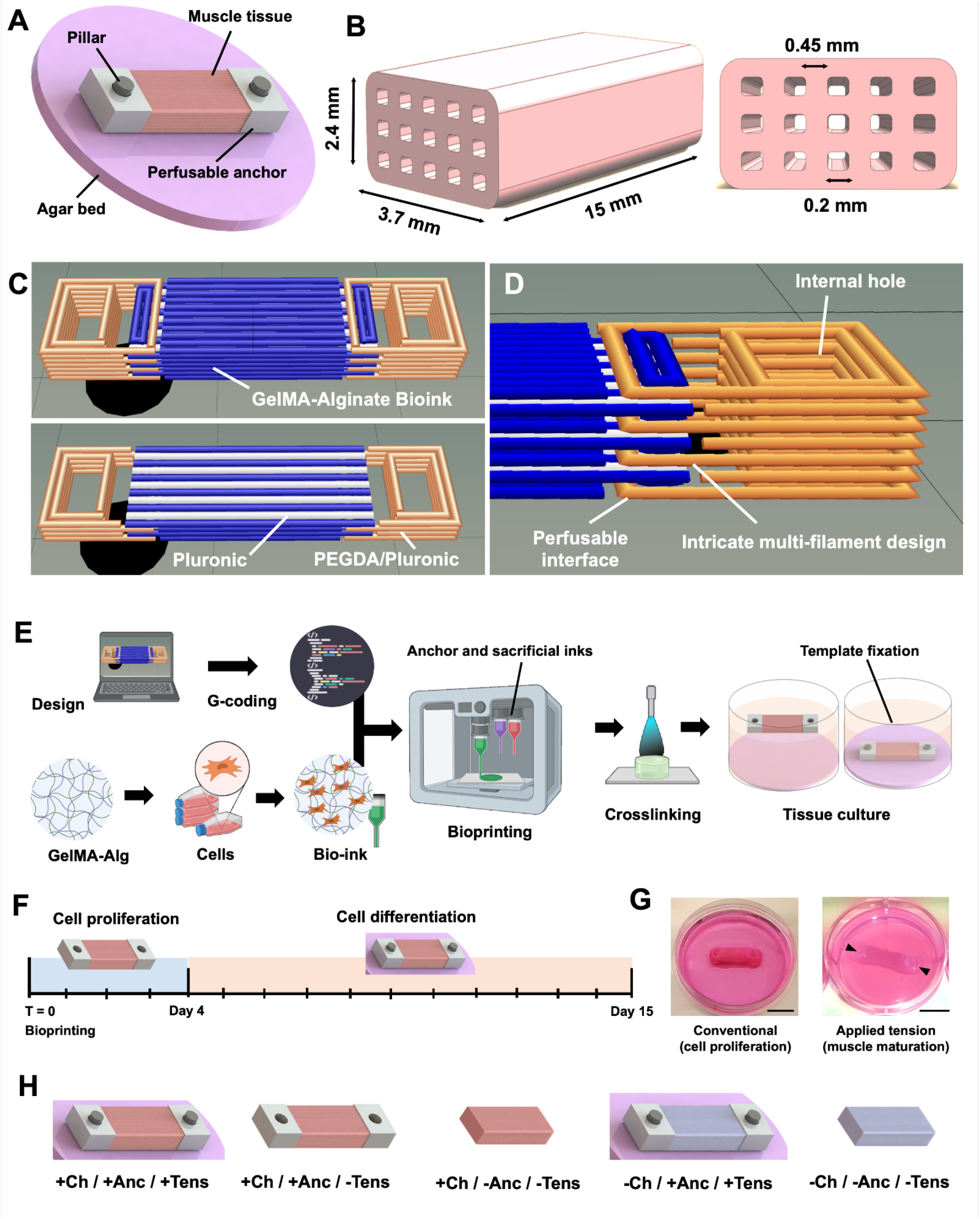
Design and realization of the biohybrid SMT. (**A**) Bioprintable design of SMT with adjacent synthetic anchors. Pillars pin down the anchors of the construct to the agar bed to form a maturation template. (**B**) Design and sizing of the internal longitudinal channels to mimic vasculature for efficient tissue perfusion. (**C**) Print design of the SMT constructs with adjacent anchor structures for all seven layers (top) and a longitudinal cut above the fourth layer (bottom). The cell-laden bioink, the sacrificial ink (Pluronic F-127), and the anchor ink (PEGDA-Pluronic F127) are shown in blue, white, and orange, respectively. (**D**) Magnified visualization of the perfusable anchor structure with hole. (**E**) Manufacturing process steps: conversion of design into G-code; selection and optimization of constituent materials; mixing with cells and formation of the bioink; extrusion-based bioprinting; crosslinking the polymeric matrix; and culture of SMT constructs under mechanical tension using a maturation template. (**F**) Culture conditions for muscle tissue maturation, including culture in growth medium for four days first, and then in differentiation medium for eleven days after mounting on the maturation template. (**G**) In the first phase of culture (left), the SMT constructs floated in the cell culture media to favor cell proliferation. In the second phase (right), the constructs were fixed through their anchors to maturation templates to receive mechanical stimulation via passive tension. (**H**) Target SMT construct and control groups used for the study (Ch = channels; Anc = anchors; Tens = tension). Scale bars: 1 cm.

The design of the biohybrid SMT constructs was optimized for high print fidelity via iterative prototyping (**Figure S1**) resulting in a multilayered, triple-material assembly with seven deposition layers (**Figure 1C**). This assembly was based on a central rectangular block fused with identical anchoring structures at the extremities. The central block (serving to generate the living tissue) contained lines of two inks, namely the cell-laden bioink (blue) and the sacrificial ink (white). These two inks were interspersed with each other and ran along the longitudinal axis of the central structure. To create perfusable channels, the lines of the sacrificial ink continued beyond the central block and interfaced directly with the external environment. The anchors were designed with an inner perfusable interface to allow for perfusion of the central block’s microchannels left upon removal of the sacrificial ink (**Figure 1D**). To stabilize the perfusable interface, the anchor ink (orange) was patterned to interweave with the bioink. The anchor ink also formed a central hole that served as the insertion point to stabilize the pillars in the maturation template (**Figure 1D**).

After bioprinting and photo-crosslinking (**Figures 1E and F**), we kept the constructs for four days in a growth medium to boost cell proliferation. We then mounted the constructs on a maturation template and cultured them in a differentiation medium for eleven days to form myofibers (**Figures 1F and G**). Constructs lacking channels (−Ch/+Anc/+Tens), anchors (+Ch/-Anc/-Tens), or both (−Ch/-Anc/-Tens) were used as control groups and cultured with the same cell culture protocol (**Figure 1H**). We also co-printed a design with channels and anchors and then let those constructs in culture without pillar-mediated fixation (+Ch/+Anc/-Tens) to see the effect of no mechanical tension during maturation (**Figure 1H**).

#### 2.1.1. Material selection and fabrication

To fabricate SMT constructs with co-integrated anchor structures, we developed a bioink with suitable mechanical properties for bioprinting, muscle tissue development, and stable co-assembly with synthetic structures. We optimized the bioink for viscoelastic behavior and printability by testing different formulations with variable relative fractions of gelatin methacrylate (GelMA) and sodium alginate (NaAlg). The fractions varied in the ranges of 4-8% and 3-11% for GelMA and NaAlg, respectively (**Table S1** and **Figure S2** to **S10**). The formulation based on 8% GelMA and 7% NaAlg displayed a linear viscosity profile over a predefined frequency range (**Figure 2A**) and a sol-gel transition depending on the applied shear stress (**Figure 2B**). Both behaviors facilitate the selection of the optimum printing parameters. Moreover, the bioink was sufficiently soft (**Figure 2B**) to support the printing and culture of myoblasts, and could be printed with an accuracy of 90% (**Figure 2C**).

**Figure 2.**
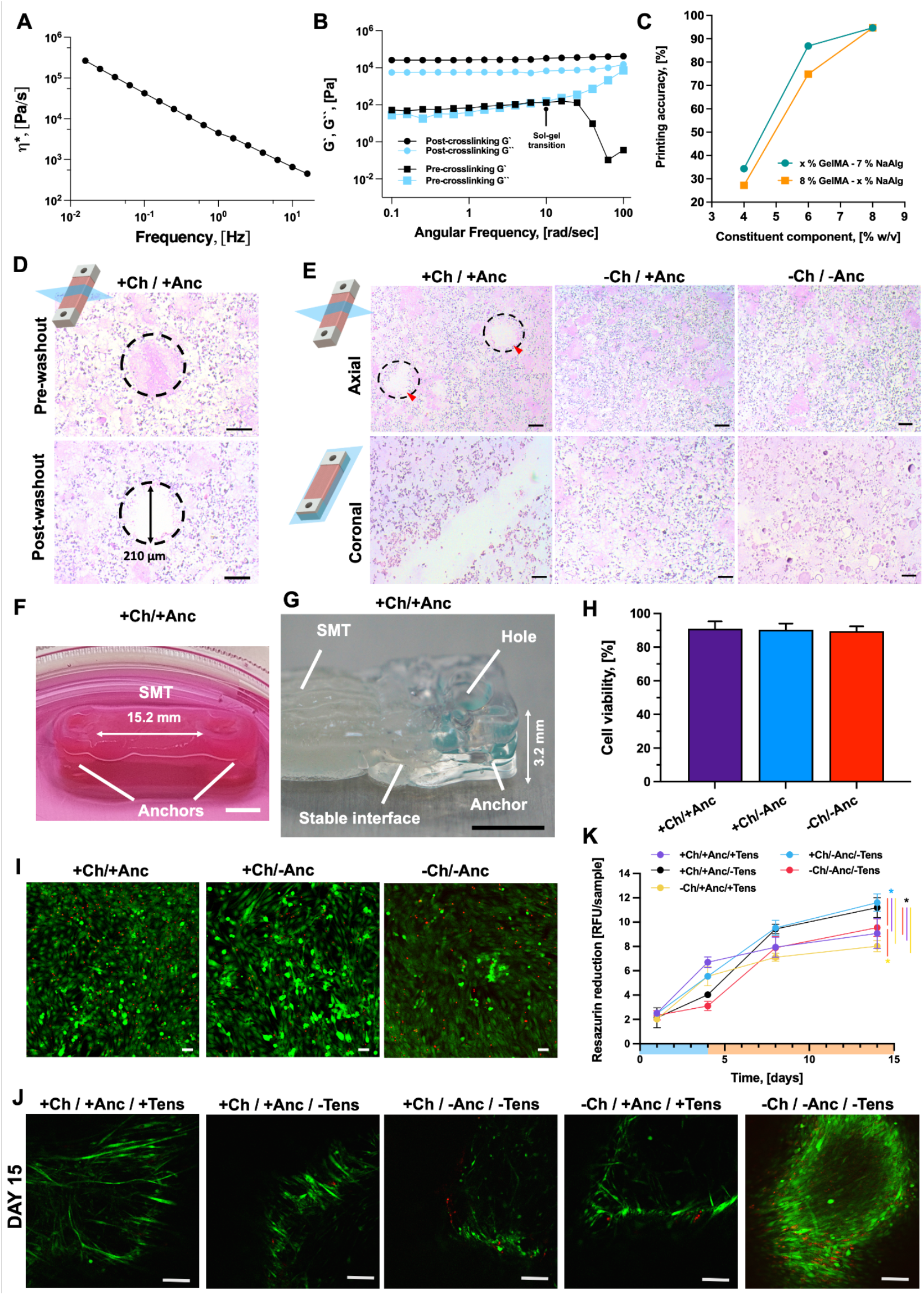
Material characterization and suitability for printing. (**A**) Variation of the viscosity (η) of the ink composed of 8% GelMA and 7% NaAlg over the shear rate (frequency). (**B**) Rheological behavior of the GelMA-NaAlg formulation before and after cross-linking. The storage (G’) and loss (G’’) moduli of the inks are plotted against the shear rate, and shown in black and cyan, respectively. Squares and circles refer to formulations analyzed before or after crosslinking, respectively. (**C**) Printing accuracy of the optimized bioink (8% GelMA and 7% NaAlg) as compared to other formulations with variable fractions of GelMA (green) or NaAlg (orange). (**D**) Histological section (H&E) of a cell-laden construct with channels showing the axial section of channels before and after wash out of the sacrificial ink. (**E**) H&E histology of SMT within the constructs after printing, as shown in the axial and coronal view (top and bottom, respectively). Red arrows and dashed lines indicate the lumen of channels. Scale bar: 150 and 100 μm (top and bottom, respectively). (**F**) Optical picture of an anchored construct prototype in cell culture media, showing the stable assembly of tissue and anchors, and (**G**) the detail of an anchor structure in a multimaterial construct prototype. Scale bars: 5 mm. (**H**) Percentage cell viability as calculated from Live/Dead staining images after bioprinting of constructs with channels and anchors (+Ch/+Anc), with channels and no anchors (+Ch/-Anc), and constructs with no channels or anchors (−Ch/-Anc) (*n* = 5). (**I**) Representative images of Live/Dead staining immediately after bioprinting. Scale bar: 150 μm. (**J**) Representative Live/Dead staining images showing cell viability and cell morphology changes at 15 days of culture. Scale bar: 50 μm. (**K**) Metabolic activity of cells over time as calculated as Resazurin reduction in relative fluorescence units (RFU) (*n* = 5). The proliferation and differentiation phases are highlighted in light blue and orange on the time axis, respectively.

The line width of the bioink and sacrificial ink was around 450 and 200 μm, respectively, and co-printing them allowed us to create architectures that mimicked the target design with high structural fidelity (**Figure S11** to **S14**). In particular, we observed (*i*) parallel microchannels running along the longitudinal axis of the constructs, which were open at the extremities and contiguous with the external space (**Figures 2E** and **S12**); (*ii*) homogeneous size of the microchannel lumen (average size: 191.7 μm ± 17.8) and interchannel distance across the construct’s volume (**Figure S13**); (*iii*) compact and homogeneous internal matrix and intact construct borders (**Figure S14**).

We generated the bioink by combining the GelMA-NaAlg blend with skeletal muscle cells (25×10^6^ cell/mL) and then co-printed the bioink with the sacrificial ink and the synthetic ink for the anchors to create biohybrid SMT constructs. Co-printing sacrificial ink with the cell-laden bioink resulted in channels with a diameter of approximately 200 μm (211± 27.3 μm), which retained their diameter after the removal of the sacrificial ink (**Figures 2D, E**, and **S13**). The biohybrid construct maintained its structural integrity after washing out of the sacrificial ink (**Figure 2F**) and the bioink and the adjacent anchor structures interfaced with each other (**Figure 2G**).

The post-printing cell viability was assessed via Live/Dead staining (**Figures 2H and I**), which showed that the cells retained high viability (> 90%) in all types of bioprinted designs. This result suggests that the presence of the inks for the channels and anchors, as well as the related bioprinting protocols, did not substantially alter the cell viability. To assess cell proliferation over time, the whole constructs were analyzed via Live/Dead staining at days 3, 6, and 15 (**Figures 2J** and **S15**). High cell viability was found in all construct designs and conditions. However, in the designs with channels (+Ch/+Anc/+Tens; +Ch/-Anc/-Tens; +Ch/+Anc/-Tens), we detected lower amounts of red (dead) cells as compared to the conditions without channels (−Ch/+Anc/+Tens; -Ch/-Anc/-Tens).

Moreover, imaging showed the effect of mechanical tension during the differentiation phase on the morphological change of cells toward myotube formation. The constructs under mechanical tension (+Ch/+Anc/+Tens; -Ch/+Anc/+Tens) displayed strong cell alignment on days 3 and 6, and long and viable myotubes at different tissue depths (100 and 500 μm) on day 15 (**Figure S15**). In the absence of mechanical stress (+Ch/-Anc/-Tens; -Ch/-Anc/-Tens; +Ch/+Anc/-Tens), the cells aligned only to a limited extent at the early maturation stage (day 6), and no evident viable myotubes were visible in the later maturation stage (day 15).

The morphological evolution of both cells and full 3D constructs complicated the imaging-based quantification of cell viability, which is why we monitored cell proliferation by quantifying cell metabolic activity with a Resazurin assay (**Figure 2K**). The constructs that were fixed to the agar template during culture (+Ch/+Anc/+Tens; -Ch/+Anc/+Tens) showed a biphasic metabolic curve that reflected the culture conditions: during the first days of culture, the resazurin reduction increased, which confirmed the proliferation of cells within the scaffold. After day 5, the rate of cell proliferation decreased, likely due to the initiation of the differentiation and the density-dependent growth inhibition. The metabolic activity of muscle tissue developed in the absence of mechanical stimulation started to decrease at a later stage (after day 8). Even though the channeled constructs were expected to have fewer cells than the ones with no channels (about 3.75×10^5^ less), their reduction in resazurin levels was always greater than those of constructs with no channels. This observation suggests that a perfusable meso-architecture can strongly affect cell viability and behavior.

#### 2.1.2. Perfusion of the scaffolds

To understand the distribution of oxygenation gradients and their effects on cell viability, we assessed the constructs for their perfusability and cell response to hypoxic gradients. Beyond 200 μm of depth, the constructs without channels displayed cell death distributed over a wide area (**Figure 3A**). In contrast, constructs with channels had fewer dying cells which were also localized within circumscribed areas, suggesting that the presence of channels supports cell viability.

**Figure 3.**
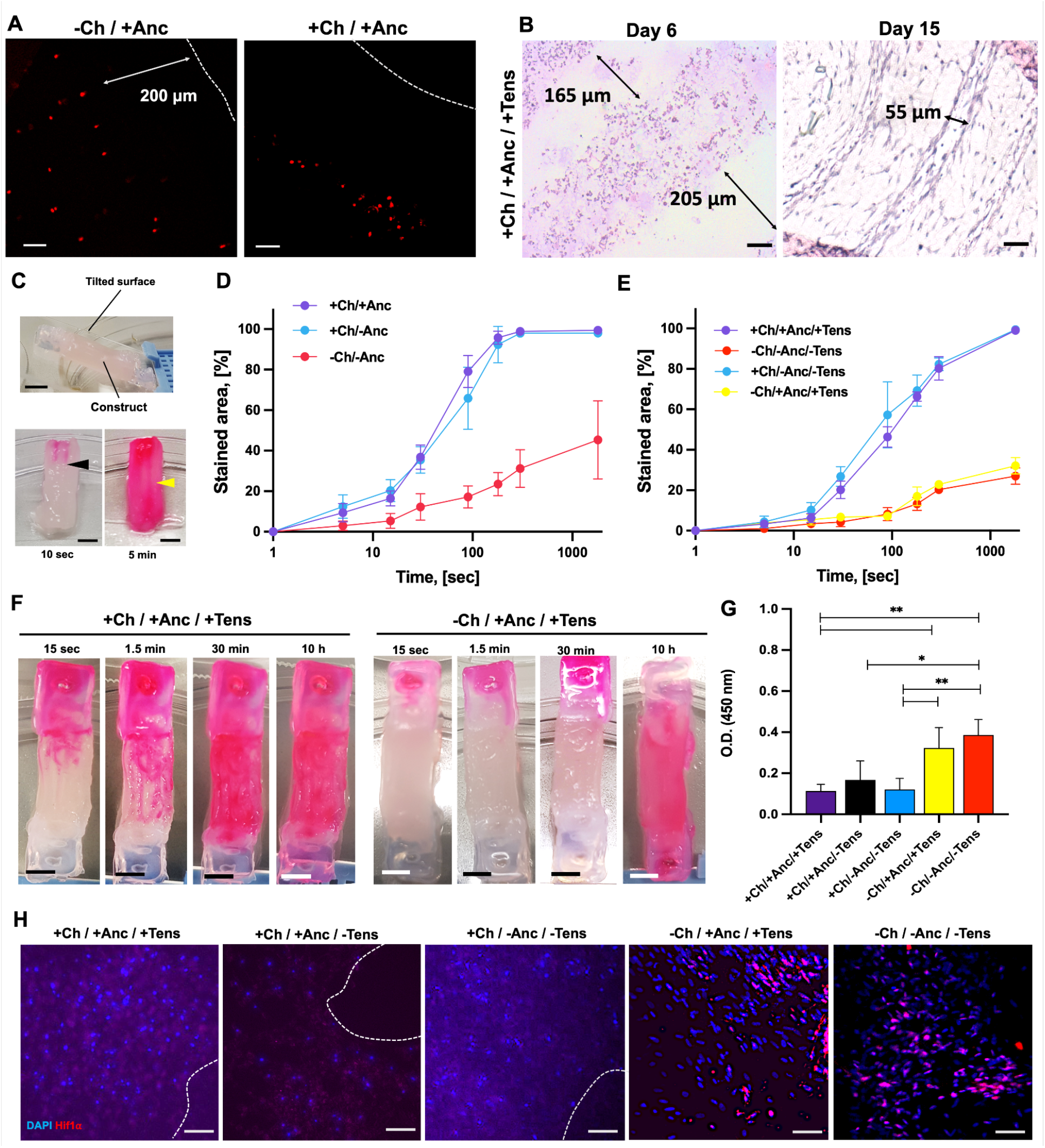
Channel perfusion. (**A**) Dead cell staining (day 3) of constructs showing cell death (red dots) at a distance of about 200 μm from the border (dashed line) of scaffolds with or without channels (left and right, respectively). (**B**) H&E of constructs at days 6 and 15 (top and bottom, respectively), as shown in coronal view. Scale bar: 100 and 50 μm, respectively. (**C**) Perfusion experiment setup with the constructs placed on tilted glass support (lateral view; top), and an added staining solution perfusing the constructs (front view; bottom). Black and yellow arrows indicate stained channels becoming clearly visible in a few seconds and remaining still evident at the endpoint of the study (5 minutes of observation). Scale bar: 3 mm. Staining kinetics reporting on the increase of the fraction of stained area over time for constructs with or without channels and anchors, shortly after bioprinting (**D**) and at the end of the maturation process (day 15) (*n* = 5) (**E**). (**F**) Optical images (frontal view) of the perfusion in the constructs at the end of the maturation process with mechanical tension (day 15). Scale bar: 3 mm. (**G**) Optical density (O.D.) at 450 nm of nuclear extracts from constructs treated for the functional assessment of the HIF-1ɑ transcription factor (*n*=5). (**H**) Confocal immunofluorescence imaging of constructs stained with the anti-HIF-1ɑ antibody (red) and DAPI (blue). Dashed lines delineate the channels’ cavities. Scale bar: 50 μm.

Coronal histological sections of the cell-laden constructs (**Figure 3B**) showed that at the end of the tissue maturation process, the size of the channels’ lumen decreased due to tension application and SMT remodeling (from 200 to 50 μm circa). However, the channels were neither collapsed nor obstructed. We verified the functionality of the channels to transport fluid and their contribution to cell viability by subjecting the printed constructs to a perfusion assay with a staining solution based on cell culture medium containing a red dye. Shortly after bioprinting, the constructs were placed in a tilted position (angle 30°) on a glass substrate (**Figure 3C**) and the staining solution was repeatedly added to one side of the construct with exposed channels. The staining solution entered the void spaces of the constructs, revealing the channel structures, and perfused the hydrogel material in a few minutes (**Figure 3C**).

By measuring over time the stained fraction of the construct from optical images, we estimated the perfusion kinetics, which revealed more than 3-fold faster staining of the channeled tissues as compared to the control groups with no channels (**Figures 3D** and **S16**). The presence of the anchors did not affect the perfusion kinetics, suggesting that the lumen of the channels was open to the external environment and the anchors did not obstruct the lumen (**Figures 3D** and **S16**). At the endpoint of the maturation process (day 15), the channeled constructs were still perfusable with faster kinetics as compared to a solid muscle without channels. These kinetics were unaffected by the presence or absence of tensioning during maturation (**Figures 3E and F**). Channeled mature biohybrid constructs were fully perfused by the staining solution after only 30 minutes of staining exposure, whereas constructs matured in the absence of channels were fully stained after 10 hours of staining exposure (**Figure 3F**).

We quantified through a colorimetric reaction the content of HIF-1ɑ transcription factor as a major regulator of hypoxia-response intracellular pathways (**Figures 3G and H**).^30^ The absence of a perfusable network within the scaffolds led to a higher amount of functional HIF-1ɑ than in the constructs with channels. In the constructs with channels developed under anchorage and tension (+Ch/+Anc/+Tens), the HIF-1ɑ levels were 3.5-fold lower than in the equivalent but not perfusable system (−Ch/+Anc/+Tens). Confocal imaging revealed co-localization of the HIF-1ɑ and nuclei signal within the non-perfusable constructs, but not in the constructs with channels (**Figure 3H**), indicating that the absence of the channels correlates with the onset of hypoxic responses in the cells.

#### 2.1.3. Tissue-anchor interface and construct stability

Despite the SMT’s deformation during maturation, the anchors retained their shape and could be penetrated by pillars for mounting to the maturation template (**Figure 4A**). The SMT and the anchor materials formed an intertwined pattern, which consolidated over time to generate a coherent interface between the two phases that was externally visible also at the end of the maturation process (**Figure 4A**, bottom). Histological analysis of the internal structure of the interface showed interpenetrating areas of the SMT and synthetic ink (**Figure 4B**, blue and red arrows). Void areas in the histological sections revealed the scheme of perfusable channels at the intersection between the tissue and the anchor (**Figure 4B**, black arrows). Confocal imaging on day 5 revealed a dense cell population on the tissue-anchor interface area and in the matrix projections within the anchors (**Figure 4C** and **S17**). In the histological sections, cells were found in the anchor areas, suggesting that cell invasion and spreading of tissue formation contributed to rendering the biohybrid interface stable and coherent (**Figure 4D**). Even if sporadic lacunae were present at the border between the SMT and anchors, the biomatrix efficiently adhered to the anchor surface (**Figure 4E**). At day 5, the surface contact between the biological tissue and the synthetic components was greater than 90%, independent of the application of tension (duration: 1 day) and the presence of channels in the SMT construct (**Figures 4E and F**). On day 15, the interface areas with full contact between the two phases decreased (remaining however >70%), suggesting that the prolonged mechanical tension decreases the stability of the assembly to a mild extent.

**Figure 4.**
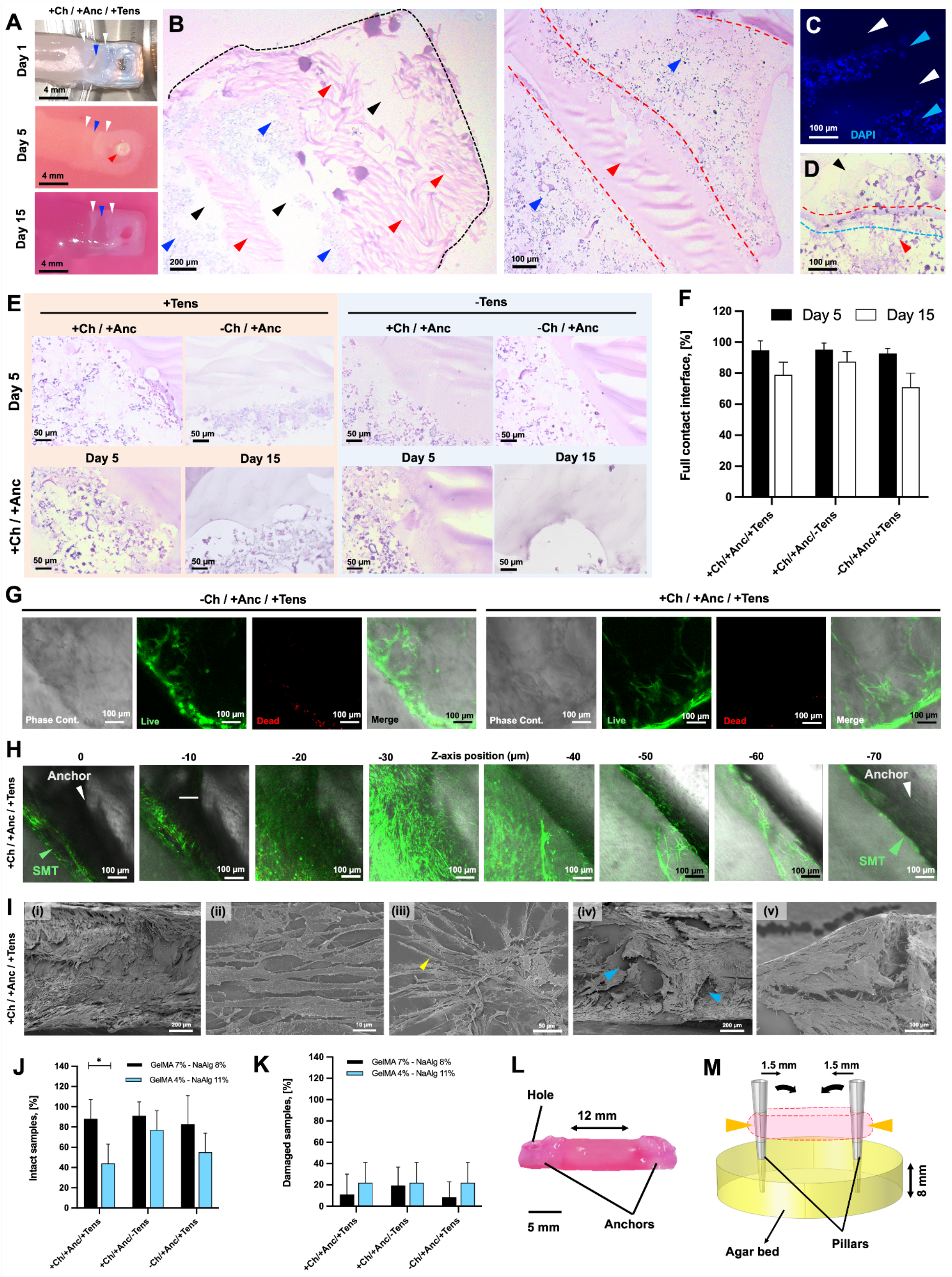
Stability of the biohybrid interface. (**A**) Photographs of the anchors with a pillar inserted in the anchor hole (red arrow), and intertwined areas of tissue and anchor structures (blue and white arrows, respectively). (**B**) H&E staining of the anchor structure and the biohybrid interface (coronal sections; day 1). Void perfusable areas, anchor material and cell-populated matrix are shown by black, red and blue arrows; the anchor appendix within the cell-laden matrix is shown by a red dashed line. (**C**) Confocal imaging of the biohybrid interface (day 5) with cells (DAPI-stained nuclei in blue, cyan arrows) growing on the synthetic surface (white arrows). H&E staining of: (**D**) the cell-laden matrix (red arrow) superimposed on the anchors (black arrow), showing two different anchor borders (red and cyan dashed lines) underneath the cell matrix; and (**E**) biohybrid interface subjected to tension or not: constructs with and without channels (day 15, top) and with channels (day 5 and 15, bottom). (**F**) Percentage fraction of full contact areas between the muscle tissue and the anchor structure calculated from histological images (*n* = 4). 3D Live/Dead staining showing viable cells on the anchor-tissue interface (**G**) and alternating layers of tissue and anchor with tissue depths (**H**). (**I**) SEM of: muscle tissue formation (i); aligned cell growth (ii); myotubes (yellow arrow, iii); cavities (blue arrows, iv) and tissue formation around them (v). Percentage fraction of intact (**J**) and damaged constructs (**K**) (*i.e*., localized structural damage with preserved functionality) at day 15 (*n* = 4). (**L**) Photograph of the perfusable construct on day 15. (**M**) Model of the maturation template with a force acting on the center of the pillars to induce their inward bending (orange arrows).

As shown by Live/Dead staining on 3D constructs, viable cells with an elongated morphology adhered and grew on the tissue-anchor interface of the constructs with channels, whereas clusters of dead and roundish cells were observed on the interface of constructs without channels (**Figure 4G**). Moreover, by exploring the anchor-tissue interface via confocal imaging along the vertical axis, we observed the alternation of the anchor structures and the tissue layers, which in constructs with channels displayed a high-density cell population with high viability index and SMT-relevant morphology (*i.e*., myotube-like cell bodies) (**Figure 4H**). As shown by Scanning Electron Microscopy (SEM), cells could adhere to the anchors’ surface while growing aligned and forming myotubes, however without invading and obstructing the designed cavities (**Figures 4I**, and **S18** to **S29**).

At the end of the tissue maturation time, the constructs were recovered for structural analysis (**Figures 4J, K**, and **S30**). Most of the constructs retained the original biohybrid conformation, with anchors stably integrated into the whole assembly (**Figure 4J**). The fraction of constructs that completed the tissue maturation while maintaining their overall integrity was approximately 90% and did not vary due to the presence of channels within the tissue or the construct fixation on the maturation template. The overall integrity was higher for the SMT based on the 7% GelMA - 8% NaAlg ink than for constructs based on a different bioink formulation (4% GelMA - 11% NaAlg). The number of constructs that got superficially damaged while remaining functional was the same for the two bioinks (<25%) (**Figure 4K**).

The structural analysis performed on the construct photographs revealed that the longitudinal size of the muscle tissue reduced from the initial value of 1.5 cm (day 0) to 1.2 ± 0.5 cm on day 15 (**Figure 4L**), and the two pillars of the maturation template converged by approximately 15° toward the middle point of the construct longitudinal axis (**Figures 4M** and **S31**). The force exerted by the muscle construct onto the pillars caused a deformation. We measured the deformation by quantifying with a cantilever system the force acting on a pillar inserted into an agar phantom (**Figure S31**). A shortening of the construct at 3 mm corresponds to a 1.5 mm displacement of each anchor; the total force required for this displacement was found to be 604.1 ± 101.5 μN, which falls in a common value range for passive forces generated by engineered skeletal muscle.^31^ These observations suggest that the selected bioink supported an SMT with appropriate mechanical properties for coherent and stable integration with the anchors and the maturation template, as well as for long-term development under mechanical tension.

### 2.2. Phenotypic characterization of the bioprinted muscle

#### 2.2.1. Muscle architecture formation

To assess the outcome of tissue maturation, the constructs were analyzed for the expression of biomolecular and morphological markers of skeletal muscle tissue after 15 days of culture. In H&E stainings, the cell nuclei of the mechanically tensioned constructs were oriented along a preferential direction (**Figure 5A**) and long myotubes (length > 150 μm) were observed (**Figure 5B**). In the constructs with channels (+Ch/+Anc/+Tens), the lumen of the channel was seen along with the cells clustered in the proximity of the channels’ walls, aligning with their axes. Not subjecting the constructs to mechanical tension resulted in matrices with a heterogeneous distribution of cells growing with multiple directionalities (**Figures 5A and C**).

**Figure 5.**
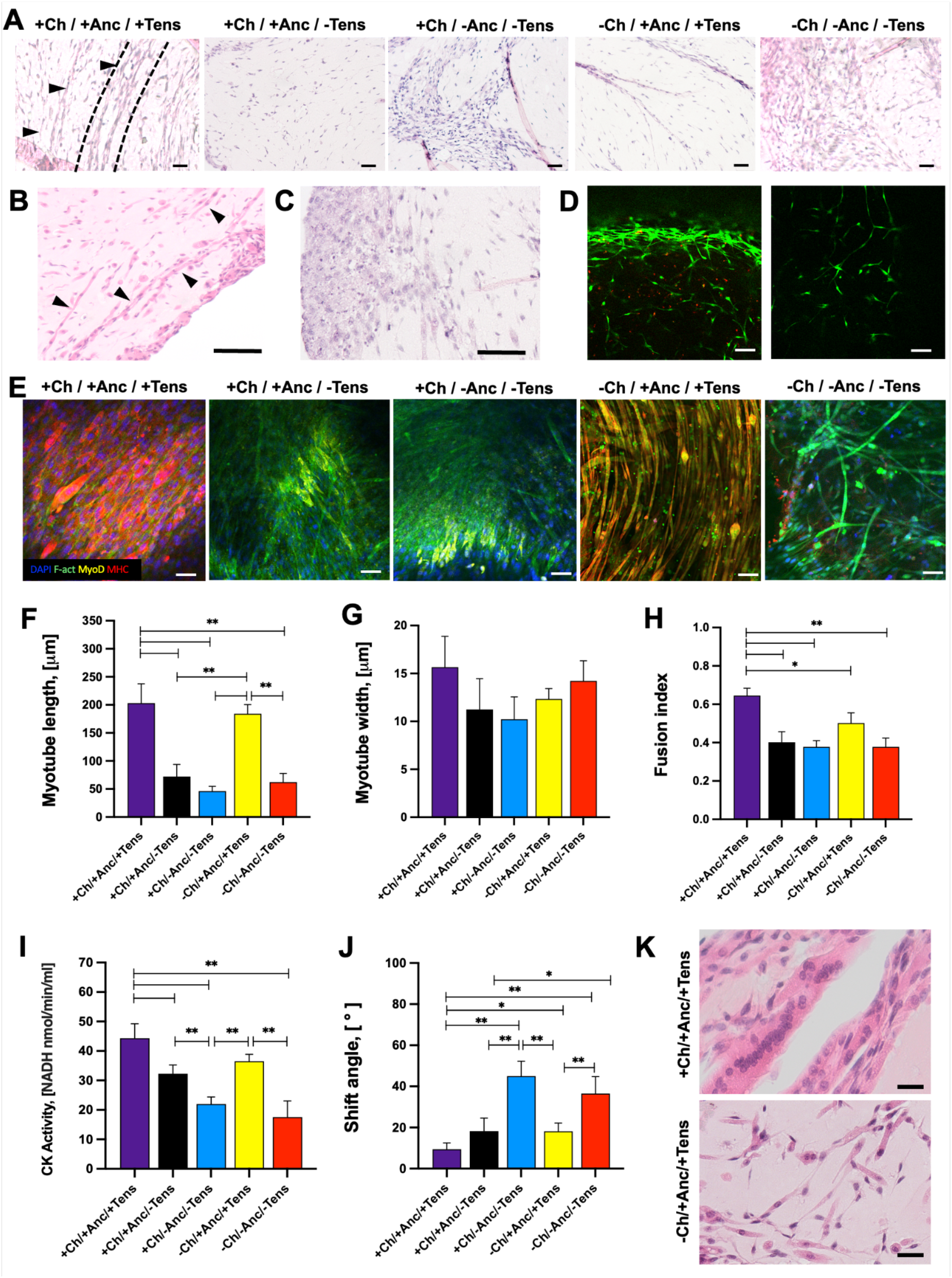
Assessment of skeletal muscle maturation. (**A**) H&E staining of constructs at day 15. Channels are visible in +Ch/+Anc/+Tens constructs (black arrows and dashed lines). Scale bar: 50 μm. **(B**) Detail of myotubes in the matrix of +Ch/+Anc/+Tens constructs (black arrows). Scale bar: 100 μm. (**C**) Heterogeneous cell distribution in the matrix of -Ch/-Anc/-Tens constructs. Scale bar: 100 μm. (**D**) Live/Dead staining of 3D constructs (+Ch/+Anc/+Tens) at day 15 on the border and internal regions (left and right, respectively). (**E**) Immunofluorescence staining of myogenic differentiation markers on the SMT constructs at day 15. Nuclei, F-actin, MyoD and MyHC are shown in blue, green, yellow and red, respectively. Scale bars: 50 μm. Myotube length (**F**) and width (**G**) as calculated from f-actin staining on confocal imaging *(n*=4). (**H**) Fusion index (*i.e*., number of nuclei inside MyHC-positive myotubes divided by the total number of nuclei present in a field of view) of constructs, calculated from confocal imaging (*n*=4). (**I**) Creatinine kinase (CK) activity as evaluated by a colorimetric assay (*n*=4). (**J**) Myotube alignment expressed as angle shift between the long axis of the myotube and the x-axis direction (*n*=4). (**K**) H&E staining shows higher cell clustering and alignment in the proximity of channels (top) than in constructs with no channels (control groups, bottom). Scale bar: 30 μm.

The expression of the major markers for skeletal muscle differentiation and maturation, namely MyoD and MyHC (Myosin Heavy Chain), varied among conditions, suggesting different stages of the muscle tissue formation process (**Figure 5E**). Tensioned biohybrid SMT displayed stronger expression of MyHC as compared to other conditions. Myotubes were observed in all conditions, but in constructs lacking mechanical stimuli they were found sporadically and with reduced size (**Figure 5F**). While myotube width did not vary significantly among conditions, the myotubes of channeled or full constructs under tension (+Ch/+Anc; -Ch/+Anc) were substantially longer (*ca*. 200 μm) than those in the other samples (ranging from 50 to 70 μm) (**Figures 5F and G**). Importantly, the fusion index and the creatinine kinase activity were significantly higher in perfusable muscle under mechanical stress than in all other conditions (**Figures 5H and I**).

Myotubes on the mechanically stimulated SMT constructs had significantly lower angle shift (15.2 ± 4.5° and 10.2 ± 6.1°) as compared with unstimulated control groups (up to 45.2 ± 7.2°) (p < 0.05), which pointed to a higher myotube alignment (**Figure 5J**). Intriguingly, in the unstimulated constructs with channels, the myotubes in the proximity of the channels were highly aligned (**Figure 5K**) with a calculated shift angle value of 18.2 ± 6.5°, exhibiting higher overall cell alignment with respect to the other control groups (**Figures 5J and K**). This observation suggests that channels can provide a structural pattern that guides cell organization and the unidirectional growth of myotubes to some extent.

#### 2.2.2. Secretome analysis

To gain deeper insights into skeletal muscle tissue maturation, the cytokine content of the culture media was analyzed via ELISA at different time points (**Figure 6**). In this analysis, several well-known regulators of myogenic functions have been considered, including four ligands for the cytokine receptors (IL-4, IL-6, IL-12, and IL-13), two ligands for receptor tyrosine kinases (BDNF and VEGF), and two members of the TGF-β family (Myostatin and BMP-4).

**Figure 6.**
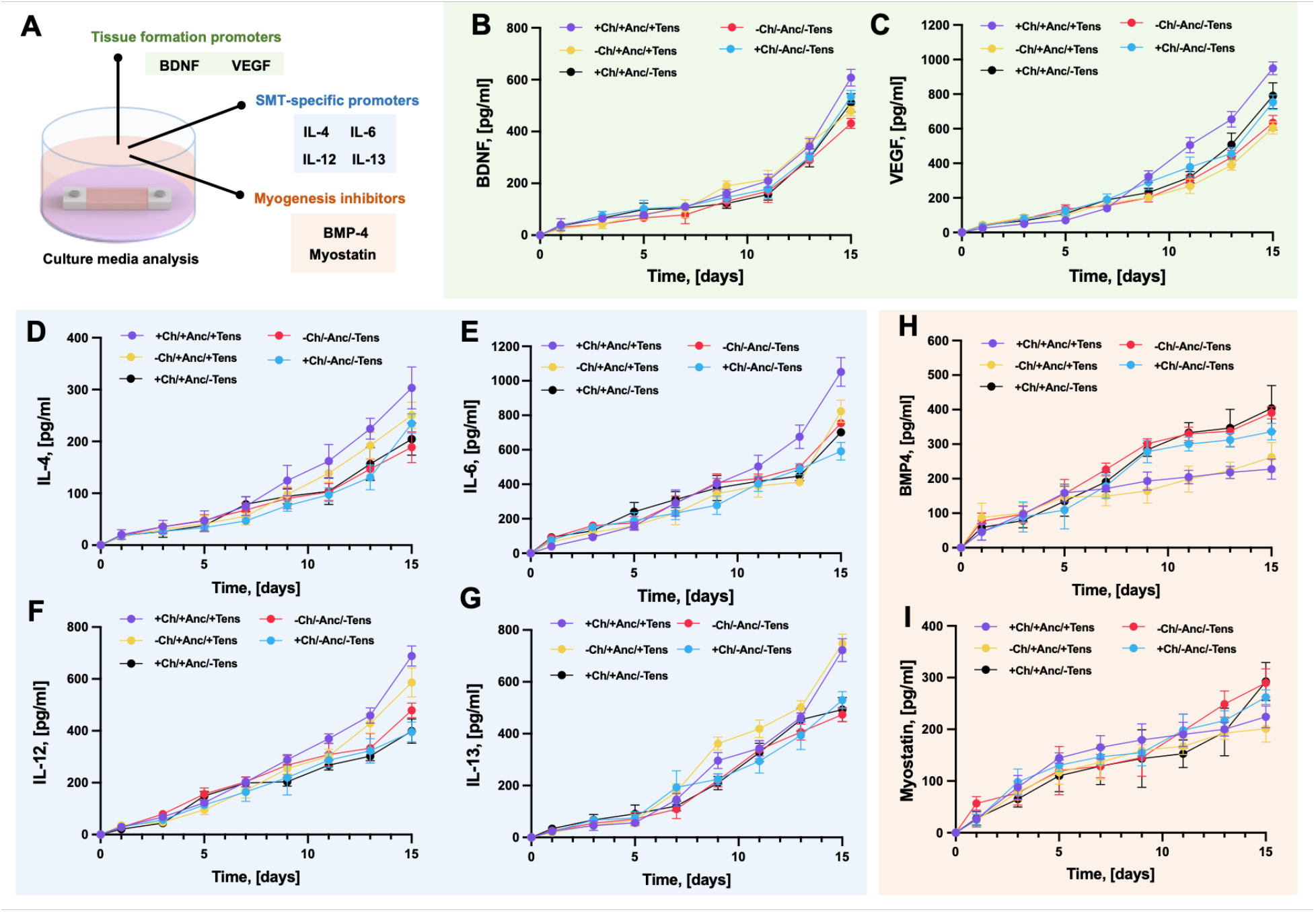
Characterization of the secretome. Schematics of experimental procedure and classification of the analyzed bio-factors (**A**). Quantification of cytokines in the culture media: BDNF (**B**), VEGF (**C**), IL-4 (**D**), IL-6 (**E**), IL-12 (**F**), IL-13 (**G**), BMP-4 (**H**), and Myostatin (**I**). The medium was collected at different time points from the culture of constructs with anchors and channels (+Ch/+Anc/+Tens); with channels and without anchors (+Ch/-Anc/-Tens); with anchors and without channels (−Ch/+Anc/+Tens); with neither channels nor anchors (−Ch/-Anc/-Tens); and constructs with anchors and channels matured without anchoring to the agar template (+Ch/+Anc/-Tens). For all assays, *n*=4.

During early myogenesis, IL-6 promotes satellite cells’ activation and proliferation. In the transition from myoblast to nascent myotube and then to mature myofiber, IL-6 stimulates cell cycle withdrawal and the initiation of differentiation, as well as cell migration and fusion.^32,33,33,34^ The highest cumulative concentration of IL-6 was found in the liquid media of constructs with channels and anchors fixed to the agar template to mature under mechanical tension (peak value at day 15: > 1 ng/mL). IL-4 and IL-13 induce second-stage fusion during myogenic differentiation.^32,35,36^ Their release was significantly augmented in constructs developed under mechanical tension as compared to the non-fixed control groups. IL-4 production was increased by more than 20% in all conditions developed under tension at all late time points (day 11, 13, and 15), while the production of IL-13 in anchored constructs developed under tension was enhanced by around 55.6 and 64.3 % at day 13 and 15, respectively, as compared to +Ch/+Anc/-Tens constructs. The +Ch/+Anc/+Tens constructs also showed increased production of VEGF, BDNF, and IL-12, which can be arguably correlated to the application of mechanical tension.

During myogenic differentiation, VEGF not only induces cell cycle withdrawal, cell migration, and differentiation, but it also promotes cell survival.^37^ The release of VEGF from cells in designs without channels was reduced as compared to the designs with channels (−36.4 and -15.8% in the anchored and non-anchored constructs, respectively, at day 15). BDNF is a recognized promoter of cell cycle withdrawal and myogenic differentiation.^32,38^ At the endpoint of the observation range, the cumulative concentration of BDNF in the media of anchored, channeled constructs reached a value of around 600 pg/mL, the highest value observed in the series of samples.

As an additional positive regulator of the differentiation initiation, IL-12 was also analyzed, showing increased production in the second part of the kinetics for all types of constructs, which points out an auto/paracrine role of this interleukin. The skeletal muscle tissue developed under tension displayed the highest values of IL-12 cumulative release peaking in the range of 500-700 pg/mL at the end of the kinetics and strongly diverging from constructs cultured in suspension. Myostatin is a negative regulator of proliferation and at the same time, it inhibits the initiation of satellite cells’ differentiation.^39^ It is expressed in diverse *in vivo* scenarios, including developing muscle, muscle regeneration following acute injury, satellite cell-dependent compensatory hypertrophy, and muscular dystrophy. In our study, all the constructs expressed myostatin in the initial phase of the culture. Two days after the beginning of the myogenic differentiation phase, the level of released myostatin decreased in all samples, with more evident inhibition effects in the constructs developed under mechanical tension. A similar trend was observed for BMP-4, a cytokine that suppresses cell cycle withdrawal and myogenic differentiation, and it is expressed in the developing muscle.^40,41^ Even if the production of BMP-4 was drastically reduced in the second phase of the tissue culture (*i.e*., differentiation phase) in all constructs, a stronger inhibition was observed in the constructs subjected to mechanical tension (+Anc/+Tens) as compared to those cultured in suspension (−Anc/-Tens; +Anc/-Tens).

## 3. Discussion

### 3.1. Biohybrid designs of SMT

SMT biofabrication should re-create perfusable anisotropic structures that mimic native tissue, such as perfusable microchannels aligned in parallel with bundles of mature myofibers. To align fibers, the maturation of engineered, mesoscale (*i.e*., cm size ranges) SMT requires constant mechanical tensioning, which can be provided by rigid culture templates counteracting the matrix shrinkage.^1,10^ Biohybrid designs for SMT maturation should contain anchoring synthetic structures to the ends of thick muscle tissue. These modular designs also facilitate the handling of fragile hydrogel-based 3D SMT constructs.^42^ Biofabrication can even model the myotendinous axis by realizing a seamless interface that optimizes force transfer from muscle to tendon.^26^ Ladd et al. modeled a myotendinous junction model by culturing myoblasts and fibroblasts on a scaffold featuring electrospun polymers that replicate the gradients in mechanical stiffness at the interface between the tendon and skeletal muscle.^43^

Despite its potential for SMT engineering, the multi-material bioprinting of synthetic 3D structures and their biohybrid interfaces has been scarcely investigated.^28^ Merceron et al. printed a myotendinous construct by alternating the deposition of polycaprolactone and polyurethane with muscle cells and fibroblasts.^44^ Both polymers were layered in a grid shape to form a backbone with two distinct elasticities that structurally support the growth of both cell types into a construct mimicking the native tissue organization. The synthetic backbone resembled a distributed scaffolding material that facilitated the integration with the cell-laden matrix. Kim et al. used multi-material bioprinting to form a cubic SMT block (one cm^3^),^28,45^ in which the bioink was co-printed with a sacrificial gelatin-based ink and synthetic support pillars but this assembly was optimized for simultaneous mechanical tensioning and communication of the channels with the external environment.

In contrast, our print design allows media to diffuse through the channels and anchors. The anchors fix the muscle to the pillars in the maturation template, and could support a longer muscle tissue block suitable for the development of long fibers. The printing pattern stably intercalated the tissue and anchor materials during the layers’ deposition. By optimizing the printing procedure against various parameters, we achieved a sufficiently high printing resolution to realize constructs with the intended characteristics, such as perfusable channels and interpenetration between SMT and anchors. Despite the shrinkage of the matrix during muscle tissue development, our biohybrid SMT remained functional for two weeks. Our design printed with PEGDA/Pluronic and GelMA/NaAlg was robust enough to withstand forces spanning the μN and mN range, which are typical for bio-actuators of similar size.^31,46,47^ Most intriguing was that patterning perfusable networks within the constructs not only reduced the occurrence of hypoxic regions without altering the structural stability of the assembly, but also provided contact guidance to the growing myocytes, promoting the formation of unidirectionally aligned muscle fibers.

In our system, synthetic materials and biomaterials were shaped into one construct during the bioprinting process. The synthetic structures were stabilized via crosslinking at the same time as the biomatrix itself. The muscle tissue then matured while being in direct contact with their surfaces, allowing the soft tissue to adapt and tightly adhere to the complex morphology of the anchors. Future studies should address if our one-go fabrication process is advantageous not only to support the SMT development but also their dynamic functions, for example by increasing the efficiency of force transmission through the tendon-like structures.

### 3.2. Materials for bioprinting and developing SMT

Materials for biofabrication should display biomechanical properties that guarantee muscle cell differentiation and proper SMT formation. Much research has been done to elucidate the role for matrix or tissue stiffness in striated muscle development,^24,25^ finding that muscle cells have complex responses to the mechanical properties of scaffolds. For example, myogenic differentiation in culture depends intimately on optimal outside-in signaling mediated by the matrix elasticity.^26^ Cell fusion into myotubes might occur independent of substrate flexibility, but myosin/actin striations only emerge in scaffolds displaying stiffness that match the values of actual muscle (passive Young’s modulus of ∼10-12 kPa).^27–30^ Unlike sarcomere formation, the cell adhesion strength augments monotonically versus substrate stiffness with the strongest adhesion on stiff scaffolds.^26^ So, for successful development of SMTs requires scaffold materials that display the right balance between elasticity and stiffness. Alginate and gelatine-derived hydrogels provide for excellent myogenic differentiation *in vitro* due to a mechanical profile that mimics one of the native matrices (*e.g*., elasticity moduli of 10 to 15 kPa).^48,50^

Our 8% GelMA - 7% NaAlg ink formulation allowed us to print construct geometries with higher accuracy compared to other formulations. Our constructs showed not only a higher fraction of functional channels with desired open extremities that remained functional for the whole culture duration, but also a more compact and homogeneously distributed bulk matrix, regular interchannel distance, and less fragile scaffold borders. Even if crossed by perfusion networks, the architecture of the whole construct was sufficiently intricate to guarantee a stable and durable interface between the muscle tissue and the synthetic elements. In this environment, we proved that the SMT could develop even if the cells were not embedded in conventional biomaterials (such as collagen, fibrinogen, and Matrigel)^45^ in which the differentiation of C2C12 cells and the formation of functional fibers is commonly achieved within one week.^16^ However, these materials are not optimal for bioprinting or larger structures, due to their cost, limited versatility, and applicability to high-resolution biofabrication. In our system, we achieved SMT maturation by allowing for more days of differentiation (11 days) after an initial phase required for cell proliferation (4 days).

We found that exposing SMT to mechanical stress within our biohybrid system increased autocrine and paracrine loops for myogenesis promotion. Bio-factors supporting tissue maturation were exponentially expressed for the duration of the culture. Mechanical tensioning and switching from growth to differentiation media lead to a decrease of BMP-4 and Myostatin in the second half of culture time, which fostered myogenic differentiation. While printing anchors did not affect the secretome, the presence of channels impacted the production of cytokines, suggesting that our biohybrid design preserved cell viability by effectively combining the channels of the soft tissue with the voids of the anchors.

### 3.3. Conclusions

Our results demonstrated that 3D extrusion-based bioprinting applies to the engineering of multi-phase constructs that include living cells, biomaterials, synthetic materials, and void volumes acting as perfusable networks. When realized through an intricate and perfusable design, this complex assembly supported the development of 3D skeletal muscle tissue, a process that entails dramatic structural deformation deriving from the shrinkage of the polymeric matrix, cell morphology changes, and cell-mediated matrix remodeling. We expect that layer-by-layer multi-material bioprinting of biohybrid SMT will impact the field of muscle tissue engineering for use in biomedicine, nutrition, and bio-robotics. In fact, we showed how this approach can overcome some of the main challenges in bioengineering of 3D muscle tissue *in vitro*, including: achieving shape fidelity in biomimetic tissue designs; retaining structural stability during tissue remodeling; providing efficient liquid exchange to core regions of constructs and avoiding hypoxic areas formation; and increasing ease of manipulation of tissue *in vitro*.^8,48,51^ As these challenges are also common for the engineering of other tissue types,^52,53^ we believe that our biohybrid fabrication approach can guide scientists in improving biofabrication also in other tissue domains.

## 4. Materials and Methods

### 4.1. Cell culture

The murine myoblast cell line C2C12 was obtained from the American Type Culture Collection (ATCC, Manassas, VA, USA) and was tested for mycoplasma (MycoAlert™ Mycoplasma Detection Kit, Lonza AG, Basel, Switzerland). Cells were cultured in monolayer at 37 °C in a 5% CO_2_-containing humidified atmosphere in complete growth medium (GM), consisting of Dulbecco’s modified Eagle’s medium (DMEM, #D6429, Sigma–Aldrich) supplemented with 10% (v/v) heat-inactivated fetal bovine serum (FBS, #F7524, Sigma–Aldrich), 2 mM glutamine, 100 U mL^−1^ penicillin, and 100 μg mL^−1^ streptomycin (all from ThermoFischer Scientific, Switzerland). Cells were grown at an initial seeding density of 5×10^3^ cells/cm^2^, and detached from flasks by trypsinization (Trypsin EDTA 0.25%, Sigma-Aldrich) at 70% of cell confluency.

### 4.2. Bioink preparation

GelMA was synthesized under minimal light exposure by coverture with aluminum foil. Briefly, gelatin was dissolved in PBS at a concentration of 10% w/v at 50 °C under constant stirring. 0.6 g methacrylic anhydride per 1 g of gelatin was slowly added to the solution, which was then kept continuously stirring for further 60 minutes. The solution was then dialyzed using a membrane with a molecular weight cut-off of 12-14 kDa. Dialysis was performed at 40 °C against a large volume of deionized (DI) water for four days, with daily water renewals. The solution was kept at 37 - 50 °C during the entire synthesis and purification to avoid thermal gelation. The dialyzed GelMA solution was diluted with PBS to a final volume of 40 mL and frozen overnight. Finally, the frozen GelMA was freeze-dried until completely dry and stored at 4 °C until used. To prepare the GelMA-NaAlg polymer blends, GelMA was used at final concentrations of 4, 6, and 8% w/v, while NaAlg was used at final concentrations that incremented by 3% while remaining in a defined range of total polymer concentration (11-15% w/v). All blends contained 0.5% w/v of LAP photoinitiator. The tested formulations are listed in **Table S1**. The blends were generated from GelMA precursor solutions at concentrations of 16, 12, and 8 % w/v obtained by diluting the freeze-dried GelMA into PBS, and adding the LAP photoinitiator. The solution was kept on a thermo-shaker at 1400 rpm and 70 °C for 30 minutes to fully dissolve the solutes, and then filtered under sterile conditions through 0.2 μm supor filters for sterilization. The alginate precursor solutions (concentrations of 6, 10, 14, 18, and 22% w/v) were obtained by gradually adding appropriate NaAlg amounts to PBS at 65 °C, under continuous stirring. The NaAlg solutions were then kept on an incu-shaker at 250 rpm and 50 °C for 8 hours, and finally sterilized by autoclaving. GelMA and NaAlg solutions were warmed in a water bath to 37 °C before bioprinting. To prepare the pluronic ink, Pluronic F-127 (Sigma–Aldrich) was gradually added to cooled PBS (4 °C) to reach a concentration of 40% w/v. The solution was centrifuged at 4 °C for 1 minute, autoclaved, and stored at 4 °C until used.

### 4.3. Rheology of polymer formulations

Rheological measurements of the polymer GelMA-NaAlg blends were performed using a rotational MCR series rheometer (Anton-Paar, AT) with a parallel plate geometry. The polymer blends were deposited on the measuring surface of the rheometer plate and the upper plate was lowered until direct contact with the material was established. Frequency sweep tests were performed with a frequency range of 0.1 - 100 rad/s and a constant shear strain of 0.5%. All measurements were performed at room temperature. All polymeric solutions were tested in both uncrosslinked and crosslinked states. For crosslinking, the material was first irradiated with a handheld UV light (UltraFire 502UV, UltraFire, Piscataway, NJ) for 15 seconds and subsequently covered with a crosslinking agent (50 mM calcium chloride, CELLINK). Extracted values were imported and analyzed in MATLAB (MathWorks, USA).

### 4.4. Design of the biohybrid SMT

All bioprinted muscle constructs were fabricated by extrusion-based bioprinting by means of a CELLINK Bio-X6 Bio (CELLINK, Boston, USA). This system, well applicable to multi-material printing, is composed of multiple dispensing modules (six different printheads), a pneumatic pressure controller, an XYZ stage/controller, and a sterile chamber for printing. The design of the multilayered, biohybrid construct was based on a G-code that facilitates the biofabrication via BIO X6 bioprinter by enabling layer-by-layer crosslinking and the technical adjustments of parameters (such as printing height and speed) during an ongoing print. The constructs were designed with 3D computer-aid design (CAD) modeling using Autodesk Fusion 360° (San Francisco, California, U.S.A.). As converted to a motion program, the CAD modeling provided the design’s paths to print the constructs with different morphologies.

### 4.5. Bioprinting of the biohybrid SMT

All printed materials (*i.e*., cell-laden and sacrificial bioinks) were aseptically loaded into different cartridges. The inks were extruded through conical nozzles with an inner diameter of 410 μm (22G) (CELLINK, Boston, USA) in a closed aseptic chamber during the printing process. Before printing, the nozzles were aligned to determine their respective X-Y-Z offsets with minimum 0.01 mm accuracy. Printing parameters were optimized against the following variables: pressure, speed, and temperature. The optimal printing conditions for each ink were determined through tests for strand assessment, printing accuracy, and shape fidelity (Supplementary Materials). **Table S2** lists the used inks with their optimized printing parameters. The parameters were used as a reference point, as their constant adjustment during printing was needed to ensure continuous ink deposition. To maximize the successful outcome of the bioprinting process, three preliminary iterations were run in the absence of cells, before the cell-laden constructs were actually bioprinted. To create the bioink, the cells were counted, and a specific number of cells were centrifuged, collected as pellets, and then mixed with a specific biopolymer volume to prepare the cell-laden bioink at a specific cell seeding density. The printing procedure was optimized to maximize the cell seeding density while retaining good mechanical properties of the resulting bioink that allowed for accurate printing. The optimal cell concentration was found to be 2.5×10^7^ cells/mL. To efficiently mix the cells and the bioink, the two bioink components were loaded into two sterile interconnected syringes, and shuffled for a minimum of 50 times (pace of 1 Hz for a minimal time duration of 50 sec). Into detail, the cells were first resuspended into GelMA, and then the NaAlg and the cell-laden GelMA solutions were mixed through the syringe shuffling. The resulting bioink was then transferred to a 3 mL printer cartridge using the Luer lock connection. To keep both the GelMA and the Pluronic F-127 in their thermally gelated state, the printbed temperature was set to 19 °C. To generate SMT constructs, the GelMA-NaAlg blends were printed with the temperature-controlled printhead and kept at 21 °C throughout the entire print. The cell-laden bioink was printed at a speed of 10 mm/s and air pressures of 30 kPa. The sacrificial bioink was printed at a speed of 8 mm/s at 75 kPa. To print the anchors, an ink based on PEGDA (M_n_ = 700; Sigma-Aldrich) and Pluronic F-127 was used. The two components were mixed in a hot environment. Briefly, in a light-protected beaker, the LAP photoinitiator dissolved in PBS at 70°C under magnetic stirring for a final concentration of 0.5% w/v. After PEGDA addition and full dissolution (40% w/v), Pluronic F-127 (40% w/v) was gradually added to the mixture. The solution was autoclaved and stored at 4°C in a light-protected tube until use. Dispensing speed and pressure of the PEGDA-Pluronic polymer blend was 75 mm/min and 780 kPa, respectively. Each layer was photocrosslinked directly after printing using the printer’s integrated UV module (365 nm, 30 seconds). After printing all seven layers, the constructs were submerged in an ice-cold crosslinking agent (50 mM CaCl_2_, 45 seconds). As Pluronic F-127 (40% w/v in deionized H_2_O) has a low critical gelation temperature (around 14°C),^1^ washing the constructs with the ice cold crosslinking agent simultaneously flushed out the sacrificial ink and crosslinked the alginate portion of the hydrogel.

### 4.6. Culture of 3D constructs

After printing, myoblast-laden constructs were let into GM for four days to allow for cell proliferation. Half volume of the culture media was collected and replaced with a new GM. At day 5, anchored constructs were mounted on their maturation templates.^2^ These templates were plastic plates with pre-formed agar beds (3 ml, 1.5% w/v) that were pre-treated with culture media to prevent nutrient depletion of the culture media by diffusion of the nutrients into the agar bed. The beds were used to fix the anchors in place during tissue maturation by pinning them down with two pipette tips. Constructs were cultured for 11 days in differentiation medium (DM), composed of DMEM/high glucose, supplemented with 2% (v/v) horse serum, 2 mM glutamine, 100 U mL^−1^ penicillin, 100 μg mL^−1^ streptomycin, 50 ng/ml IGF-1, and 2 mg/ml 6-aminocaproic acid (ACA) (all purchased from Sigma–Aldrich). Control constructs with no anchors were simply cultured in DM without any mechanical fixation. During culture, half media content was daily renewed and aliquots of the media were collected for protein content analysis at specific time points.

### 4.7. Perfusion experiments

To assess perfusion, a dynamic staining experiment was performed. The constructs were put on a tilted glass surface (30 ° C) and three aliquots of red staining solution were repeatedly added to the upper extremity of the construct (time interval among additions: 1.5 minutes). The staining solution was composed of culture medium containing the red dye Erythrosin B (0.1% w/v; Sigma-Aldrich; product number: 200964). The staining solution was left to perfuse or diffuse tissue constructs following gravity. The stained (%) fraction of the construct was calculated from optical pictures at different time points to describe the perfusion kinetics. An additional time point for construct observation was added at 10 hours to illustrate the staining saturation conditions of all constructs.

### 4.8. Live/Dead staining in 3D constructs

To assess cell viability in the constructs, Live/Dead™ staining was performed immediately after bioprinting and at different time points during the culture time, following the manufacturer’s instructions (Thermo Fisher Scientific; product number: R37601). Calcein and propidium iodide were excited at 488 and 561 nm laser wavelengths, respectively, and imaged on a confocal microscope (Zeiss LSM 780 Airyscan, Zeiss AxioObserver.Z1). `Imaging analysis were performed on at least 3 images from each analysed sample feature (≥ 3 samples/condition; ≥ 3 experimental replicates)`.

### 4.9. Immuno-fluorescence in 3D constructs

After *in vitro* culture, the constructs were fixed overnight in a 4% Paraformaldehyde solution at room temperature. After fixation, samples were rinsed in PBS (5 minutes, 3×), and stained for f-actin, MyoD, and Myosin Heavy Chain (MyHC). Briefly, samples were permeabilized with 0.1% Triton X-100 (20 minutes) and blocked with a 1% Bovine Serum Albumin (BSA) solution (1 hour). After extensive washing, the constructs were stained with the primary Anti-MyoD Antibody (#ZRB1452, Sigma-Aldrich, dilution 1:1000) and incubated for 4 hours with an anti-rabbit secondary antibody coupled with Rhodamine (#SAB3700846 Sigma-Aldrich, dilution 1:200). Cells were incubated in AlexaFluor 488 phalloidin (#R37110, Invitrogen) for one hour. 4′-6-diamidino-2-phenylindol (DAPI) was finally used to label the nuclei (dilution 1:1000; 15 minutes). All incubations were performed under gentle shaking and at room temperature. To reveal HIF-1a, the constructs were stained with anti-HIF-1a antibody (ab179483, Abcam, Cambridge, U.K.), which was used at 1:50 dilution with goat anti-rabbit IgG (Alexa Fluor® 488) secondary antibody. The staining of MyHC was performed by incubating the samples for 4 hours with the mouse anti-MyHC antibody (Myosin 4, eFluor™ 660, Clone: MF20, Affymetrix eBioscience™), which was used at a 1:10 dilution for a final concentration of 10 μg/mL. All images were taken using a confocal microscope (Zeiss LSM 780 Airyscan, Zeiss AxioObserver.Z1, ScopeM), using the same exposure time, and were analyzed in Fiji ImageJ.^3,4^ Differentiated myotubes in a specific microscopic field were observed under ×10 and ×20 magnification. Either the total number of nuclei or the number of nuclei within MyHC-positive myotubes was counted in 5 fields/sample. The fusion index was calculated as a ratio between the number of nuclei within MyHC-stained myotubes and the total number of nuclei. The images were randomly selected from at least 3 regions from each sample, and the experiments were replicated three times. The alignment of myotubes was evaluated by measuring the angle shift between the long axis of the myotube and the y-axis direction (corresponding to the longitudinal axis of the constructs) by using ImageJ.^5^

### 4.10. Histology

After *in vitro* culture, the constructs were fixed overnight in a 4% Paraformaldehyde solution at room temperature. Then, the samples were embedded in paraffin and sections of 4.5 μm thickness were cut with a microtome (Microm, HM430, Thermo Scientific). The sections were stained with Hematoxylin-Eosin (#GHS116 and #HT110116, respectively from Sigma Aldrich). The slides were mounted, and imaged in bright field with a light microscope (Olympus CKX41, Olympus Schweiz AG).

### 4.11. Scanning electron microscopy

The samples were fixed with 2.5% (v/v) glutaraldehyde in PBS solution at room temperature for 30 min and then rinsed three times with PBS. Then, the samples were dehydrated in an ascending series of ethanol solutions (30%, 50%, 70%, 90%, and 100% (v/v); 5 min in each solution), followed by 3 incubationa for 10 min each in 100% ethanol, dried over molecular sieve. After transfer into metal capsules, the samples were inserted into a critical point dryer (tousimis 931) and the ethanol was substituted against liquid CO_2_. Then the samples were dried over the critical point of CO2 (31°C/73,8 bar). After the pressure was slowly released, the samples were taken out and mounted on SEM-stubs. For conductivity the samples were sputter coated with 5nm Pt/Pd (Safematic CCU-010). The examination was done in a JSM-7100F JEOL SEM at 3kV by secondary electron detection.

### 4.12. Metabolic activity in 3D constructs

To evaluate cell growth within the scaffolds, the AlamarBlue assay was performed to determine the metabolic activity of seeded cells and confirm their proliferation. Briefly, the culture medium was supplemented with 10% (v/v) of AlamarBlue reagent solution at 0.1 mg mL^−1^ (#R7017, Sigma-Aldrich) and incubation was carried out for 2 hours at 37 °C before the fluorescence signal (Ex/Em = 530/590) was measured with a Synergy H1 microplate reader (Biotek). Fluorescence intensity values were corrected for the background control (culture medium with resazurin).

### 4.13. Measurement of Creatine Kinase (CK) activity

The Creatine Kinase (CK) activity was used as a marker of myogenic differentiation.^6^ To assess CK activity in the constructs, a small amount of tissue (5 mg) was dissected from the constructs at the end of maturation time (day 15) and tested via a colorimetric CK Activity Assay Kit (Ab155901, Abcam, Cambridge, UK) according to the manufacturer’s instructions. The tissue was washed with cold PBS, resuspended in 100 μL of ice cold CK Assay Buffer, and homogenized with 10 passes. Insoluble materials were removed by centrifugation (5 minutes, 2000 rpm, at 4 °C). Supernatants were collected and plated at different dilutions into a 96 well plate. The Reaction Mix was added into each standard and sample well, and Background Reaction Mix was added to the Background control sample well. The output in optical density (OD) at 450 nm was measured on a microplate reader in kinetic mode for 10 minutes at 37 °C. CK activity was measured 2 times at 5 minutes intervals and each assay was performed in duplicate. The amount of NADH generated by CK during the reaction time (ΔT) was calculated from the OD as follows:

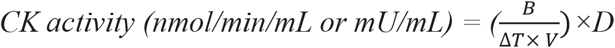

where B = Amount of NADH in sample well calculated from standard curve (nmol); T = reaction time (minutes); V = original sample volume added into the reaction well (mL); D = sample dilution factor.

### 4.14. Secretome analysis

The amount of IL-4, IL-6, IL-12, IL-13, BDNF, VEGF, BMP-4, and Myostatin in the liquid media of constructs was quantified by specific ELISA, according to the manufacturer’s protocol. Briefly, media aliquots were collected at different time points and added to an ELISA kit plate coated with an antibody against the protein of interest (capture antibody), then a biotinylated secondary antibody (detection antibody) was added and the plate was incubated at room temperature for 2 hours. The reaction catalyzed by horseradish peroxidase was stopped by adding 1 M sulfuric acid and absorbance was measured at 450 nm. Mouse IL-4, IL-6, and IL-13 ELISA kits were purchased from R&D system (Minneapolis, MN, USA; product numbers: M4000B; M6000B; M1300CB). IL-12 and VEGF ELISA kits were purchased from Thermo-Fisher Scientific (Waltham, MA, USA; product number: BMS616). BMP-4, Myostatin, and BDNF ELISA kits were purchased from LS Bio (Seattle WA, USA; product numbers: LS-F23502; LS-F35789; LS-F23504).

### 4.15. Replication of construct deformation

The passive force exerted by the muscle constructs on the pillars during tissue development was estimated via a custom-made cantilever system. The culture template composed of an 8mm-tick layer of agar gel (1.5% w/v agarose, Sigma-Aldrich) and filled-in P10 pipette tips was built in a petri dish (**Figure S31**). The dish was then fixed in a vertical position to orient the pillars horizontally. Pressure was applied to central part of the tip to achieve a later displacement of the pillar of about 1.5 mm (which was observed on real constructs). The whole set-up was placed on a scale, and weight values were taken before and after the application of pressure. The force value was estimated from the weight difference between the two conditions.

### 4.16. Data analysis and statistics

Data were analyzed with Graph Pad Prism 9 Software. All variables are expressed as mean ± standard deviation (SD). Data have been acquired from at least four independent experiments and three technical replicates, unless otherwise stated. Samples (*i.e*., bioprinted constructs) were at least three per category. To assess statistically relevant differences between the two experimental groups, the *t*-test was used (*p* < 0.05 and *p* < 0.01 are expressed as * and **, respectively). A general linear two-way ANOVA test was used to verify the hypothesis of whether there were changes in various parameters over time among the experimental groups and to identify relevant variations among several experimental groups. For all experiments, at least three samples were used to study each condition, and at least three experimental replicates were performed.

## Acknowledgments

We acknowledge Stephan-Daniel Gravert for contributing to the rendering of the schematics of muscle constructs. We also thank Anne Greet Bittermann from the Scientific Center for Optical and Electron Microscopy (ScopeM) of ETH Zurich for acquiring images at the Scanning Electron Microscopy.

## Author contributions

MF and RK conceived the original idea.

MF designed the study; performed experiments relating to material characterization, cell biology, biomolecular assays; performed analysis; wrote the manuscript.

OY performed rheology and synthesis of GelMA.

JG performed the bioprinting and material characterization.

RG designed muscle actuators and optimized the bioprinting.

AB performed COMSOL simulation of the muscle shrinkage.

RK conceived the study; came up with the idea of building biohyrid muscles.

MF, OY, RK revised the manuscript.

All authors approved the final draft of the manuscript.

