## Supplementary Material for "Perfusable biohybrid designs for bioprinted skeletal muscle tissue"

6. Perfusion of the constructs………………………………………………………. 19
7. Interface characterization ……………………………………………………….. 20
8. Stability of the biohybrid constructs…………………………………………….. 26
9. Final remarks…………………………………………………………………….. 27
10. References ………………………………………………………………………. 32
1. **Design for SMT with adjacent anchors**

The design of the anchored SMT constructs herein presented were optimized across various modeling-fabrication iterations. **Figure S1** displays an initial construct design as visualized by Repetierhost. The constructs were created as assemblies of multiple deposited layers (**Figure S1A**). Seven layers were used to realize these structures. In the pictures, the blue filaments represent the cell-laden bioink portion, while the white and orange filaments represent the sacrificial Pluronic F-127 ink and anchor ink (composed of PEGDA 700). The G-code was written using absolute coordinate commands to enable the use of layer-by-layer crosslinking and adjustment of printing parameters (e.g., height and speed) during an ongoing print. The anchors were designed as open structures to allow perfusion of the microchannels (**Figure S1B**). This design was based on an interwoven scheme of different filaments, in which single filaments of the anchor ink and the bioink were supposed to be deposited parallely to each other. The result would have been the penetration of the synthetic ink within the muscle tissue along the longitudinal axis of the construct. Such a design would result in a stabilized biphasic structure only when combining materials with highly compatible physico-chemical properties, which can generate an adherent interface. The bioink and the anchor materials were not similar from the point of view of the composition, and stabilized molecular interactions between them were unlikely to occur. Moreover, this design was too intricate to be efficiently printed with our setup.

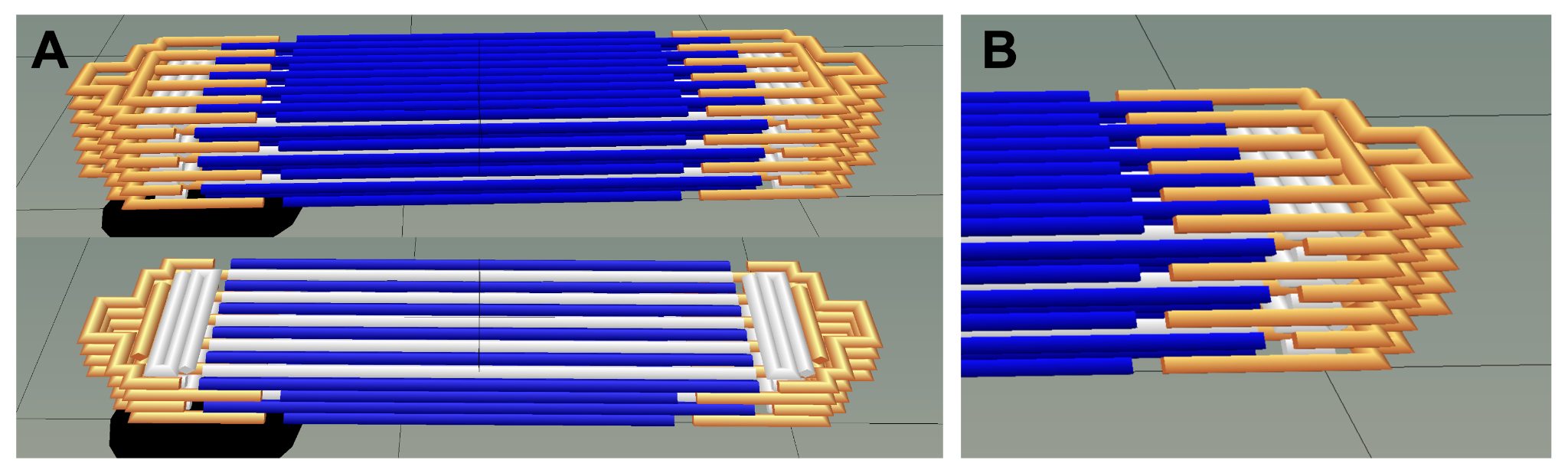
**Figure S1. Initial design for the SMT with adjacent anchor structures visualized by Repetierhost.** (**A**) The whole seven-layered construct (top) and a longitudinal cut above the fourth layer (bottom) are shown. The cell-laden bioink, sacrificial ink, and anchor ink are shown in blue, white, and orange, respectively. (**B**) Detailed structure of the anchor-tissue interface with interpenetrating inks.

The initial design was optimized in terms of architecture and materials for realization. The structure of the anchors was changed to the final design shown in the main text. Briefly, the anchor ink interfaced the muscle tissue in a way to include perpendicular structures in the scheme of assembly of the two inks. Moreover, the structure of the hole and the scheme of deposition were simplified so as to match the limit of spatial resolution that could be achieved in our printing setup. Finally, the material for the anchor filament was changed from pure PEGDA 700 to a mixture of PEGDA and Pluronic to enable printing and a more coherent interface with the cell-laden ink. In summary, as the initial designs for the whole constructs and the sole anchors were found too intricate for printing in the present setup (**Figure S1**), the architecture of the anchors was simplified so as to require a minor space resolution for realization.

2. **Characterization and optimization of the bioink formulation for bioprinting**

**Synopsys:** to generate a suitable bioink, various compositions of GelMA and NaAlg were prepared and characterized (**Table S1**).

| **Formulation** | **GelMA (%, w/v)** | **Alginate (%, w/v)** | **Total polymer (%, w/v)** |
| --- | --- | --- | --- |
| **1** | 4 | 7 | 11 |
| **2** | 4 | 9 | 13 |
| **3** | 4 | 11 | 15 |
| **4** | 6 | 5 | 11 |
| **5** | 6 | 7 | 13 |
| **6** | 6 | 9 | 15 |
| **7** | 8 | 3 | 11 |
| **8** | 8 | 5 | 13 |
| **9** | 8 | 7 | 15 |

**Table S1.** Bioink formulations tested for printability with variable fractions of GelMA and NaAlg components.

The different formulations were rheologically characterized in both crosslinked and non-crosslinked states, and their storage and loss moduli (G' and G’’, respectively) were determined over a wide shear stress range (**Figure S2** and **S3**). All the crosslinked samples exhibited a shear-thinning behavior, evidenced by decreasing viscosity at increasing shear rates. In fact, the G' values fell below the increasing G'', suggesting that all blends acquired a more liquid-like behavior at higher shear rates (**Figure S3**). Such a shear-thinning behavior is desirable, as it enables the material extrusion at low-pressure ranges (between 40 and 80 kPa), which do not hamper cell viability. In the formulations containing 6% GelMA or 7% NaAlg, augmenting the NaAlg or GelMA resulted in a coherent increase of the complex viscosity (**Figure S4**). In addition to exhibiting a shear thinning behavior, all the tested formulations were printable (**Table S2**). The 8% GelMA - 7% NaAlg formulation displayed the highest printability as shown on the strand assessment, accuracy, and shape fidelity tests (**Figure S4-S10**). The total polymer concentration of the bioink blend was the primary determinant for printing accuracy (**Figure 2C**). The formulations with overall polymer concentrations approximating 15% w/v displayed a high printing accuracy (>90 %), whereas 13 and 11% w/v polymeric concentrations caused the printing accuracy to decrease to approximately 60 and 50%, respectively. The lowest accuracy values were observed in the formulations containing lowest GelMA amounts (4% w/v), suggesting that GelMA is more critical for printing accuracy and shape maintenance than NaAlg.

**2.1 Rheology**

The various formulations of GelMA and NaAlg that were investigated in the present study for potential use as bionks underwent a rheological characterization (**Figures S2** and **S3**). Rheological tests were performed using a rotational MCR series rheometer (Anton-Paar, AT) with a parallel plate geometry. The bioink samples were deposited on the measuring surface of the rheometer plate using a 3 ml syringe. The upper plate was lowered until it contacted the boink, and the space between the parallel plates was entirely filled with material. Frequency sweep tests were performed with a frequency range of 0.1 - 100 rad/s and a constant shear strain of 0.5%. All measurements were performed at room temperature. All bioinks were tested in both an uncrosslinked and crosslinked state (as shown in **Figures S2** and **S3**, respectively). For crosslinking, the bioink was first irradiated with a handheld UV light for 15 s and subsequently covered with crosslinking agent (50 mM calcium chloride, CELLINK). Additionally, one temperature ramp was performed by increasing the temperature from 14°C to 35°C. Extracted values were imported and analyzed in MATLAB (MathWorks, USA).

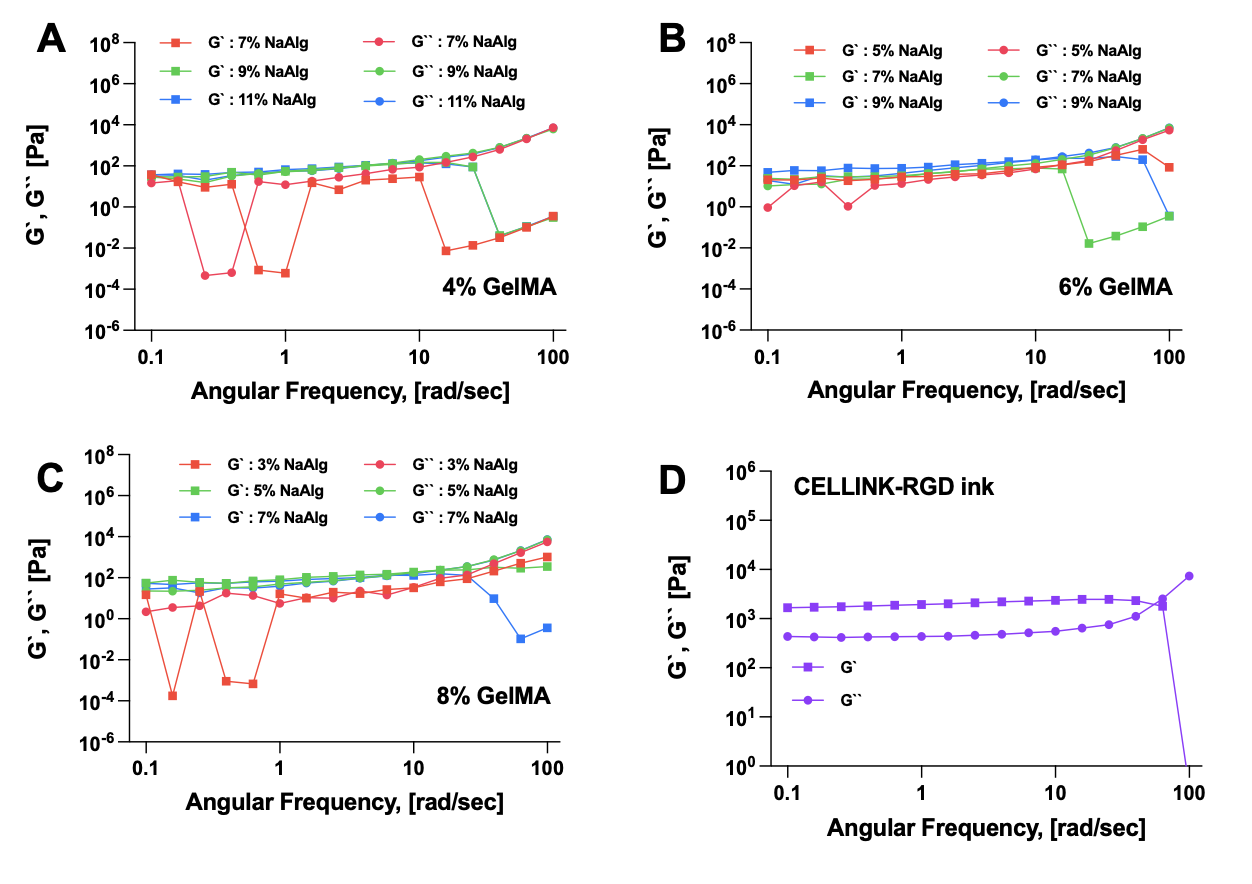

**Figure S2. Rheological behavior of the GelMA-NaAlg formulations before cross-linking.** The diagrams show the storage (G’) and loss (G’’) modulus of the composite inks as plotted against the shear rate. Storage and loss moduli are shown in squares and circles, respectively. The colors code for the different formulations obtained by increasing the NaAlg concentration (from 3 to 11% w/v) within the GelMA content fixed as 4 (**A**), 6 (**B**) and 8 (**C**) %. The rheological behavior of a commercial ink based on nanocellulolse (CELLINK-RGD, CELLINK) is reported as a comparative example (**D**).

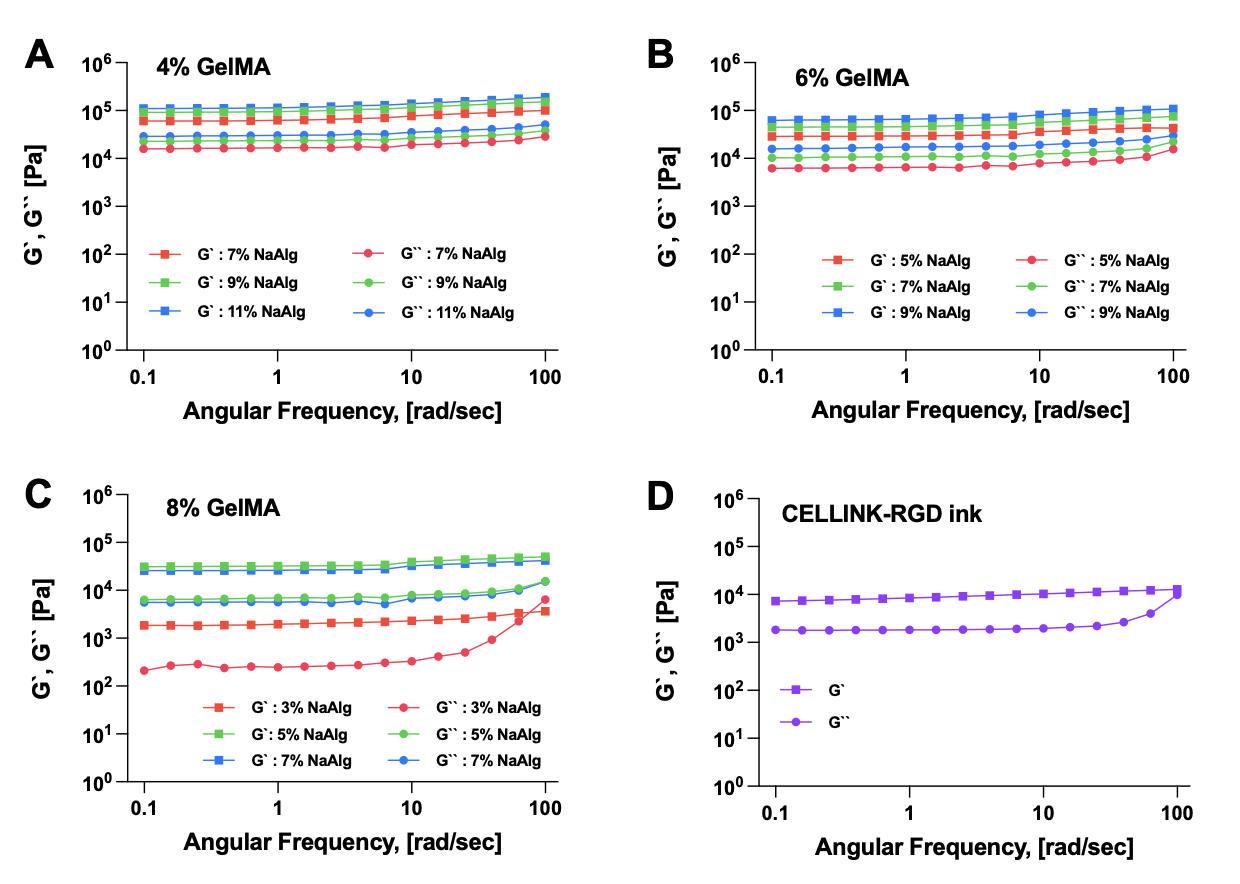

**Figure S3. Rheological behaviour of the GelMA-NaAlg formulations after cross-linking.** The diagrams show the storage (G’) and loss (G’’) modulus of the composite inks as plotted against the shear rate. Storage and loss moduli are shown in squares and circles, respectively. The colors code for the different formulations obtained by increasing the NaAlg concentration (from 3 to 11% w/v) within the GelMA content fixed as 4 (**A**), 6 (**B**) and 8 (**C**) %. The rheological behavior of a commercial ink based on nanocellulolse (CELLINK-RGD, CELLINK) is reported as a comparative example (**D**).

The formulations displayed a shear-thinning behavior as viscosities decreased with increasing shear rates. The elastic modulus (G’) fell below the increasing loss modulus (G’’), suggesting that all blends exhibit a more liquid-like behavior at high shear rates (**Figure S2**). This shear-thinning behavior is a very favorable characteristic for bioinks since it allows for material extrusion at biologically acceptable low-pressure ranges. The complex viscosities of the formulations containing GelMA at 6% or NaAlg at 7% are evident in **Figure S4**. By augmenting the Alginate or GelMA content, an increase of the complex viscosity is coherently obtained.

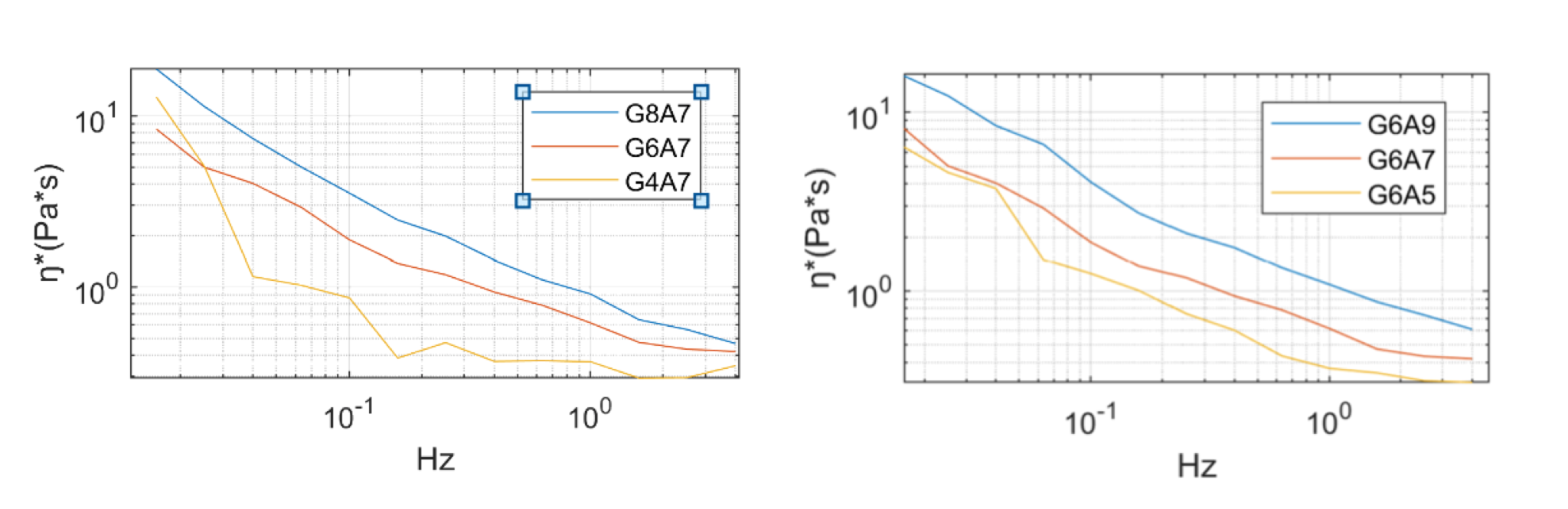
**Figure S4. Viscosity of the ink formulations.** Variation of ink viscosity (η) with increasing concentration of GelMA (left) and NaAlg (right). Legend: G=GelMA; A=NaAlg; numbers indicate the percentage composition.

3. **Bioprinting optimization**

The printing parameters were first optimized through variable value ranges for pressure, speed, and temperature. The optimal printing conditions for each ink were determined through extrudability, accuracy, and shape fidelity tests. First, the extrudability of the bioinks was first evaluated by filling a cartridge with 1 ml of material and trying for extrusion in mid-air. In this assay, the printing pressure was increased from zero by 1 kPa until the first continuous extrusion of material was observable. Then, strand assessment and printing parameter refinement were performed as follows. A continuous serpentine line was designed through Repetier Host, a 3D Printing software used for the creation of the design and setting of the printer constraints (**Figure S5**). The serpentine line was directly rendered from G-code and did not have to be sliced. The design was 5 mm in width and 10mm in length. The print height was set to 0.4 mm. Multiple serpentine lines were printed, each at different pressures and print speeds. The tested pressure ranges varied slightly between the bioinks. In general, the lower limit was set as the pressure at which continuous extrusion was achieved, whereas the upper limit was set at pressures at which an excessive amount of material started to be extruded. Print speeds of 4-14 mm/s with an increment of 2mm/s were tested. By varying speed and pressure, an overall amount of 281 combinations were tested. The conditions in which no continuous extrusion was achieved or excessive material deposition was observed were excluded from further tests. Images were taken with macro-lens on a digital 16-megapixel camera and analyzed with ImageJ (NIH, USA). The diameter of each line was measured at three different locations and averaged.

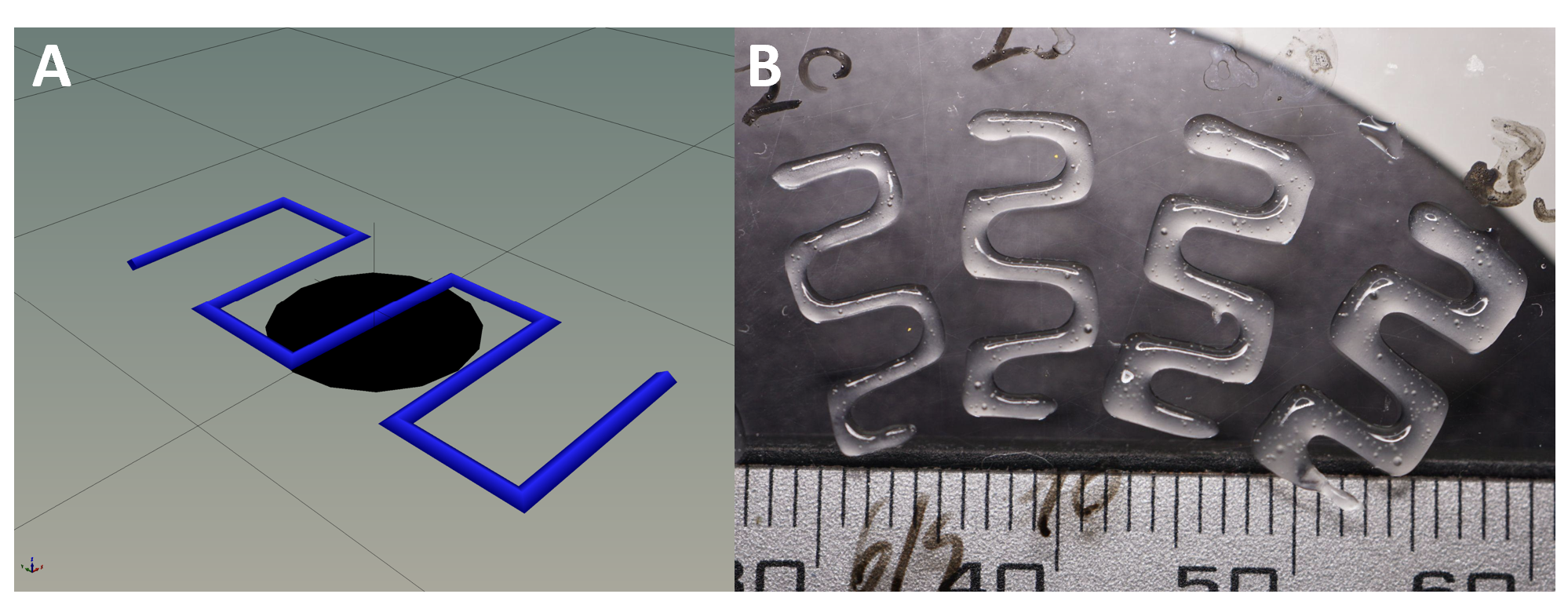
**Figure S5. Strand assessment and refinement of printing parameters.** (**A**) Visualized G-code for a serpentine line with the dimensions of 5 mm width and 10 mm length. (**B**) Assessment of the 6% GelMA - 9% NaAlg bioink. Four serpentine lines were printed at different pressures with a printing speed of 10 mm/s. The test was repeated for several different printing speeds and all nine bioinks.

All the tested GellMA-NaAlg formulations were extrudable from pressures of 10-20 kPa upwards, as expected for formulations with total polymer concentration ranging between 11 and 15%.^1^ The printing parameters impact the resulting strand width of the print. All line widths were significantly larger than the inner nozzle diameter. This is due to the Barus effect, which describes the swelling of viscoelastic pseudoplastic fluids when exiting a small orifice.^2,3^ **Figure S6** shows the relationship between printing speed, pressure, and strand width for the formulation consisting of 8% GelMA and 3% NaAlg. The printing speed has a more significant effect on strand width at high pressures, and the relationship displays an exponential trend for high pressures, while it resembles a nearly linear trend at low-pressure values.

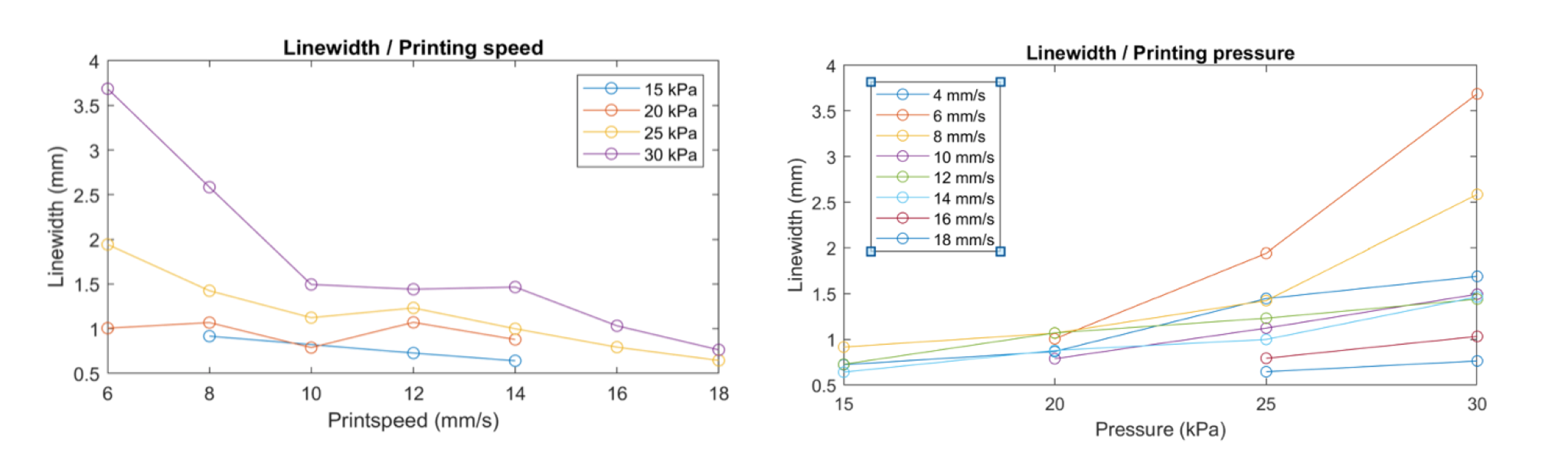
**Figure S6. Influence of printing speed and pressure on the line width.** The relationship becomes exponential at high pressures. The graphed data is exemplary and stems from the 8% GelMA - 3% NaAlg composition.

The minimal extrudable strand thickness decreased while increasing the NaAlg and the GelMA concentration (**Figure S7**). The smallest strands were observed with the blends of 8% GelMA and 7% NaAlg, with a strand width of 0.447 mm. This value is very close to the theoretical resolution, which is determined by the inner nozzle diameter of 0.41 mm.

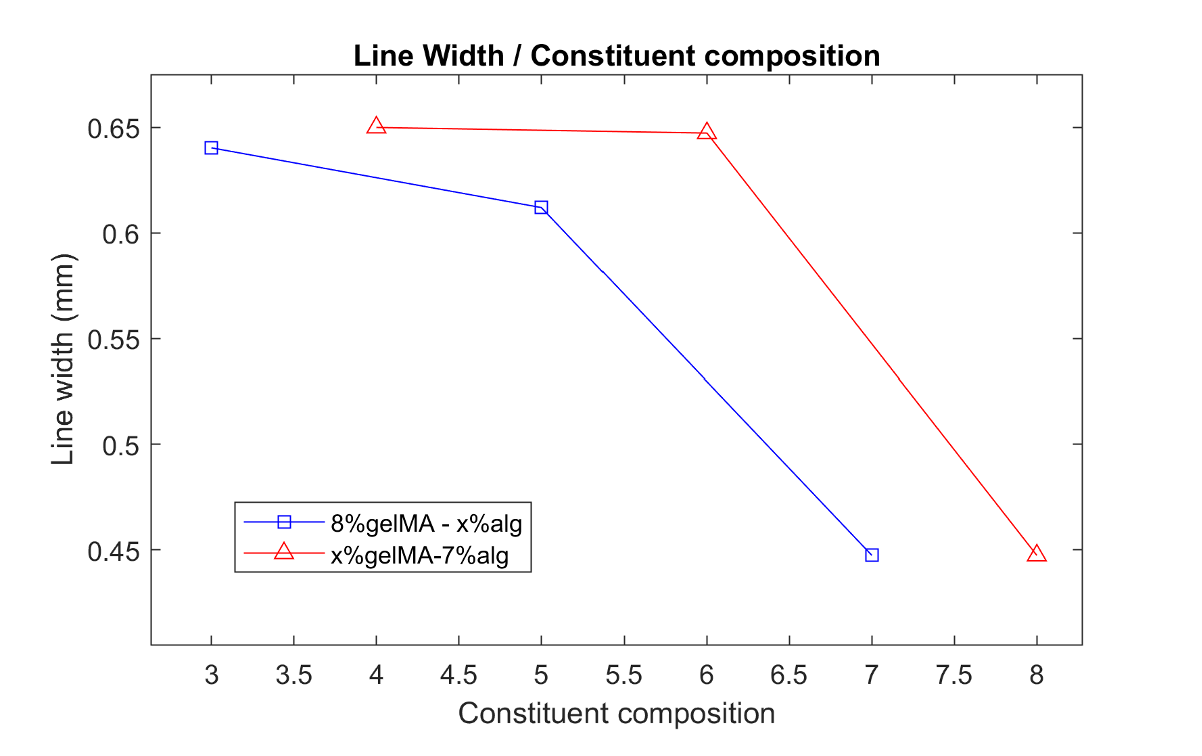
**Figure S7. Line width variation in relation to different bionk compositions.** Decreasing minimal line width with increasing concentrations of GelMA and NaAlg.

To optimize extrusion-based bioprinting, it is desirable to achieve high printing accuracy while also retaining high cell viability. The printing accuracy is inversely related to the strand width, a parameter that was extracted by analyzing the pictures obtained from the experiment described above. The cell viability depends on several factors involved in the extended bioprinting procedure. Among these factors, the shear stress represents a significant and controllable variable, as it majorly depends on the applied pressure during printing and the selected nozzle diameter.^4^ Therefore, The Parameter optimization index (POI) was calculated for each print with the following formula:

POI = $\frac{1}{t_{line}\times D_{G}\times\rho}$

where $t_{line}$ is the line thickness in mm, $D_{G}$ is the nozzle gauge (22G = 0.41mm), and $\rho$ is the printing pressure in kPa. The calculated POI values were normalized relative to the maximum POI of a specific bioink.

$${POI}_{n,i} = {POI}_{i}\div{POI}_{max}$$

Where *i* denotes the POI for an individual set of parameters while *max* denotes the maximum POI of the entire range tested and *n* is the number of discrete parameter combinations tested.^4^ The maximum normalized POI correlates to the set of parameters that maximize accuracy and minimize theoretical shear stress.

The POI was calculated for each set of parameters and normalized over the maximum POI of the relative bioink (**Figure S8**).

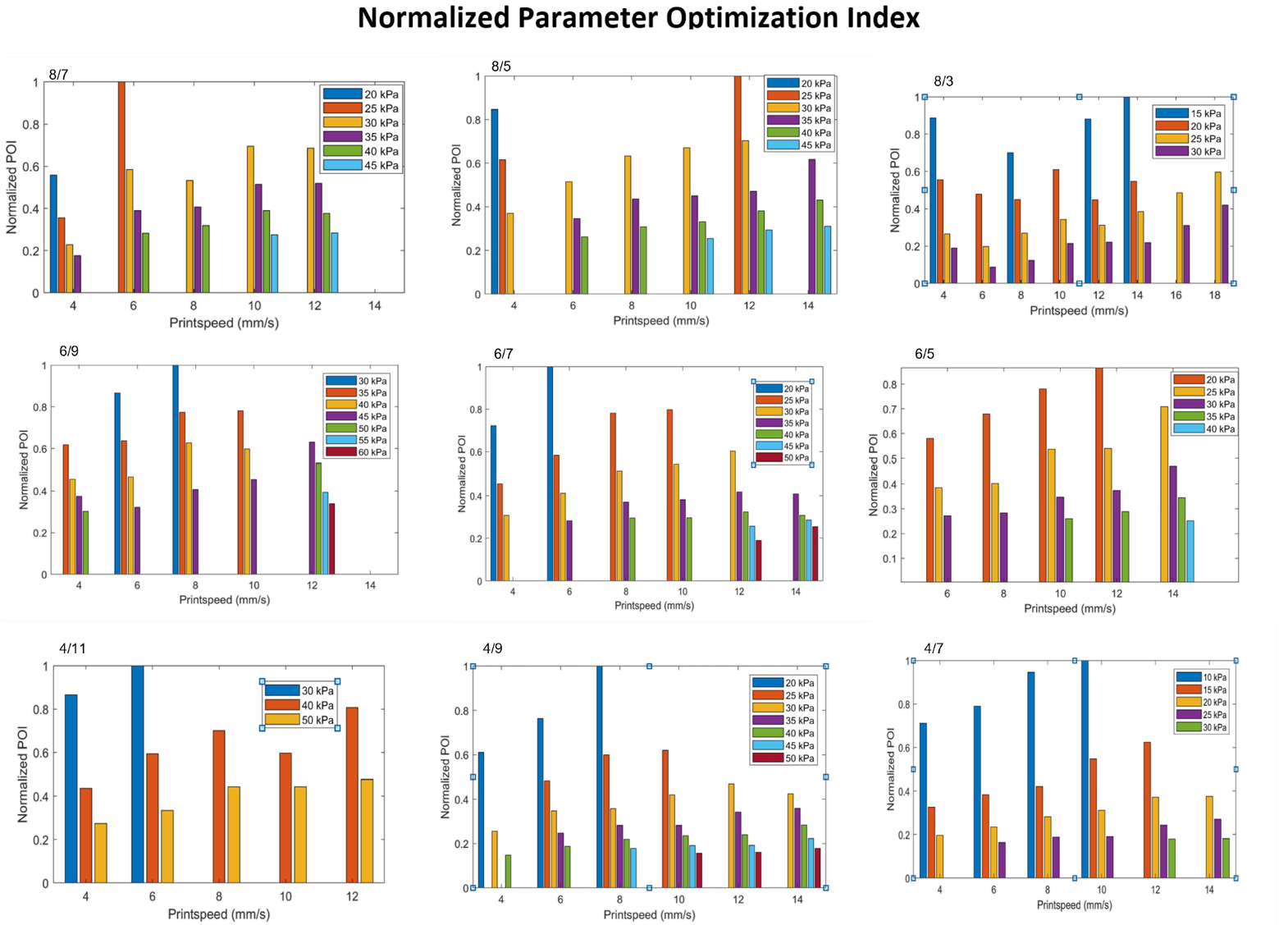

**Figure S8. Parameter optimization indices (POIs) for various combinations of printing speed and pressure.** The POI values are reported for all tested bioink formulations at variable printing speeds (x axis) and pressure values (color coding). The numbers at the top left corner of the graphs indicate the percentage composition of the constituents GelMA and NaAlg, respectively.

The maximum POI values of the different bioink blends are summarized in **Table S2**.

| **Optimizing Printing Parameters** | | | |
| --- | --- | --- | --- |
|  | **Printing pressure** | **Printing speed** | **Printing Temperature** |
| 8% GelMA - 7% NaAlg | 30 kPa | 10 mm/s | 18 °C |
| 8% GelMA - 5% NaAlg | 25 kPa | 12 mm/s | 18 °C |
| 8% GelMA - 3% NaAlg | 15 kPa | 14 mm/s | 18 °C |
| 6% GelMA - 9% NaAlg | 30 kPa | 8 mm/s | 18 °C |
| 6% GelMA - 7% NaAlg | 25 kPa | 10 mm/s | 18 °C |
| 6% GelMA - 5% NaAlg | 20 kPa | 12 mm/s | 18 °C |
| 4% GelMA - 11% NaAlg | 30 kPa | 6 mm/s | 18 °C |
| 4% GelMA - 9% NaAlg | 20 kPa | 8 mm/s | 18 °C |
| 4% GelMA - 7% NaAlg | 10 kPa | 10 mm/s | 18 °C |
| Pluronic F-127 | 75 kPa | 8 mm/s | Room Temperature |
| PEGDA - Pluronic F-127 | 120 kPa | 6 mm/s | Room Temperature |

**Table S2. Optimized printing parameters for printing materials.** The optimal printing pressure and speed for extruding different tested materials are reported for 18 °C and room temperature, through 22G nozzles.

Interestingly, the highest POI was achieved with the 4% GelMA and 7% NaAlg blend, which has one the lowest combined polymer concentrations overall. This result is due to the fact that a continuous extrusion of this blend was achieved at a very low pressure of 10 kPa, and the ink still maintained a limited line width (around 0.65 mm, see **Figure S7**).

To test the printing accuracy, a single-layered 3 × 3 grid was designed in 3D Builder (Microsoft, USA) and sliced via Slic3r to create the necessary G-code. The grid had dimensions of 10 × 10 mm and six evenly spaced 2 × 2 mm holes (**Figure S9)**. In general, due to the pooling of the extrude at the nozzle tip, the strand diameter tends to be greater than the internal nozzle diameter. As such, the strand diameters that were previously measured with optimal printing parameters were used for the Nozzle diameter input as required by Slic3r. Nine different G-codes were hence created for the following assessment. Using the optimized parameters, the grids were printed for each bioink. Images were obtained as described before, and the printed area A was measured for each grid with ImageJ. The acquired values were then compared to the design area of 64 (mm^2^). The percentage printing accuracy was obtained via the following formula:

Printing accuracy (%) = [$1 - \frac{\left| Ai - A \right|}{A}] \times100$

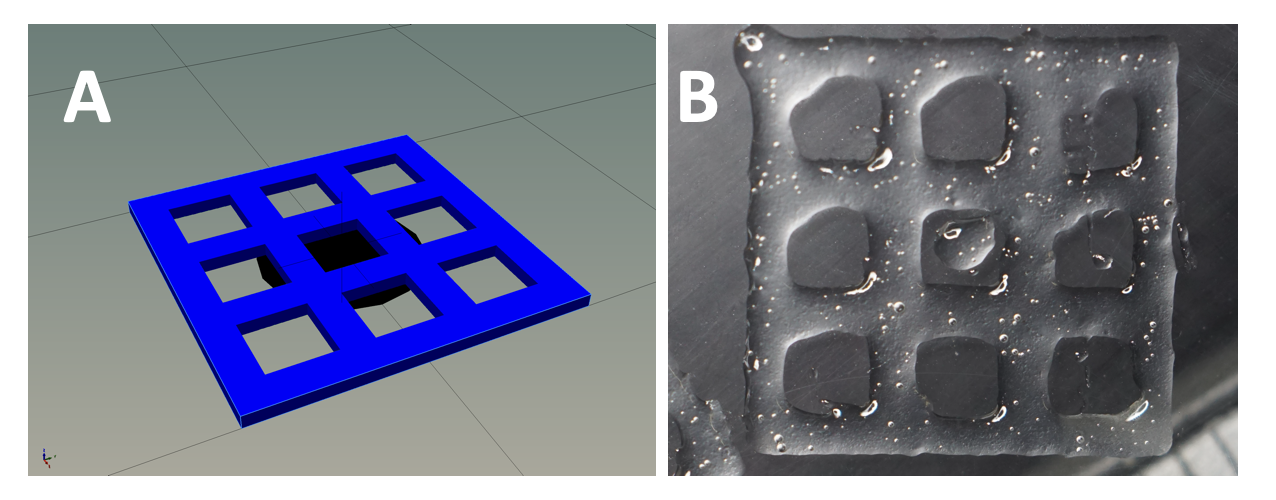

**Figure S9. Printing accuracy test.** (**A**) CAD design of the 3 × 3 grid and (**B**) 3D printed 3 × 3 grid.

The overall polymer concentration of the bioink blend is the primary determinant for printing accuracy. The formulations with overall polymer concentrations approximating 15% w/v displayed a high printing accuracy (>90\%), except for the 4% GelMA and 7% alginate. The printing accuracy for polymer concentrations of 13% w/v exceeded 60%, whereas for the less concentrated formulations (11%), it was limited. The minimum accuracy values were observed in the formulations containing the smallest GelMA amounts (4% w/v), suggesting that GelMA plays a critical role in printing accuracy and shape maintenance with respect to Alginate (**Figure S9**). Figure 2 (in the main manuscript) reports on the variation of the printing accuracy based on the constituent composition. The percentage printing accuracy (%) augmented with increasing concentrations of GelMA and Alginate. Extremely low printing accuracies were calculated with deficient GelMA concentrations.

To test the shape fidelity, a cylinder with inner and outer diameters of 8 and 10 mm, respectively, and a height of 5 mm was designed in a 3D builder, and then the design was sliced with Slic3er setting the layer height to 0.41 mm. The structures were printed using the optimized parameters (**Table S2**). Pictures were obtained as before, and the height of the printed construct was measured using ImageJ.

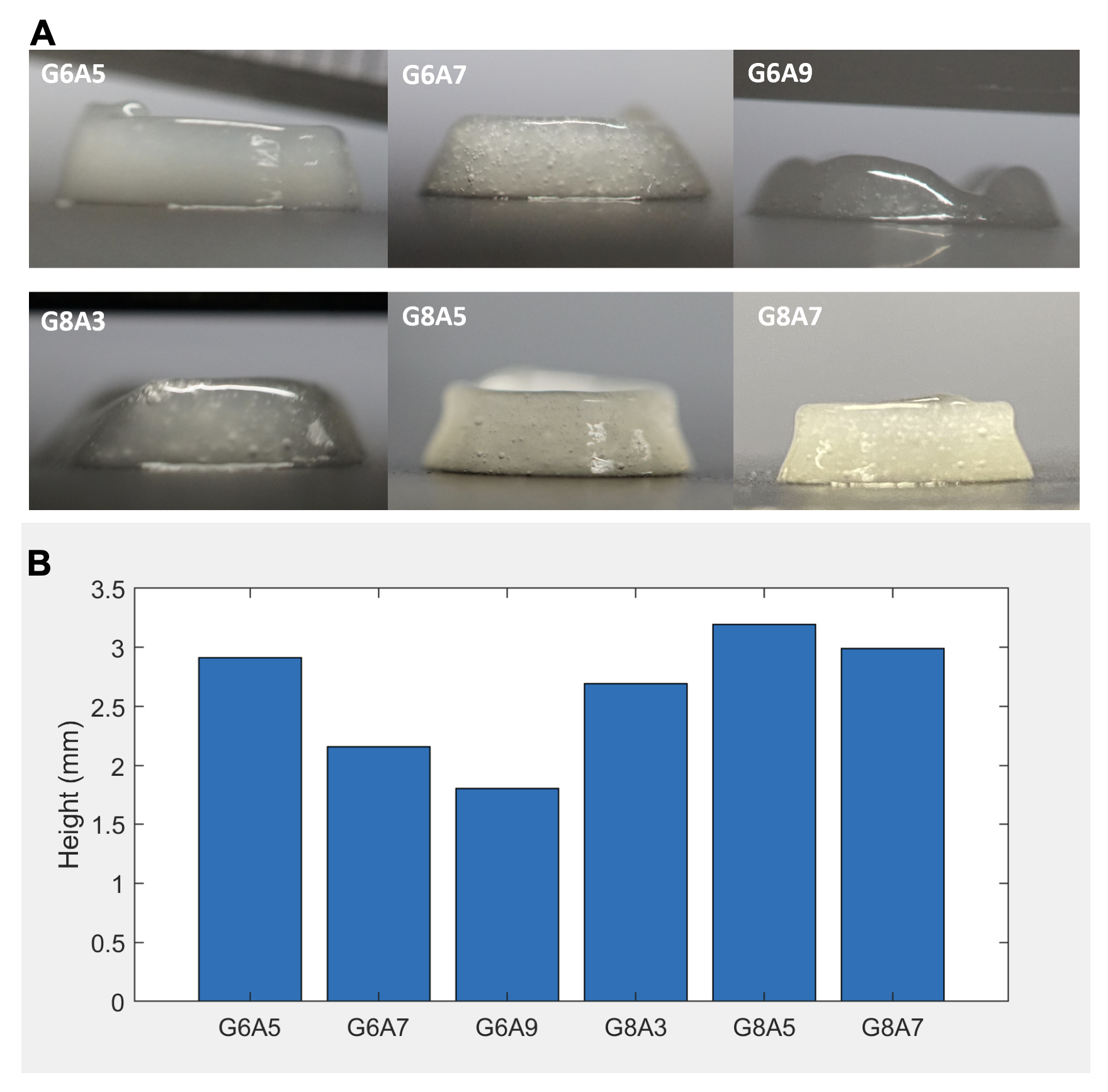
**Figure S10. Shape fidelity test.** (**A**) Optical photographs of hollow cylindrical structures printed for testing shape fidelity in bioprinting. Only few formulation produced intact structures. GelMA was photocrosslinked right after the deposition of each layer. (**B**) The height of the cylinder samples as measured from images. G = GelMA; A = Alginate. Numbers indicate percentage composition of the constituent.

The GelMA-Alginate bioink could not support a multilayered structure in the uncrosslinked state. As such, the GelMA component of the bioink was photocrosslinked after every printed layer. This way, a 10-layer cylinder was achieved by using the bioink blends containing 6 and 8% GelMA (**Figure S10A**), whereas the blends with 4% GelMA could not support ten layers and collapsed. Increasing the GelMA concentration resulted in enhancing shape fidelity. Conversely, augmenting the alginate decreased shape fidelity and resulted in collapsing structures. For all compositions, the biomaterial expanded at the bottom of cylinders due to the weight of the added layers on top. A decrease in the sample height occurred when adding alginate to the 6% GelMA formulation, whereas in the compositions containing 8% GelMA such a relationship was not observed (**Figure S10B**). Overall, these results are consistent with the notion that GelMA concentration acts as the primary determinant for the elastic modulus of the bioink, and through the addition of alginate an increase in loss modulus and therefore flowing behavior is introduced.^5^ Furthermore, as only the GelMA portion of the bioink was photocrosslinked during printing, the constructs containing lower amounts of GelMA (in combination with the uncrosslinked alginate) had a high probability to collapse while additional layers were deposited. These results indicate that to keep appropriate shape fidelity a relatively high gelMA concentration is necessary. In this experiment, the results were strongly influenced by minimal adjustments of the pressure and printing speed, which were necessary to complete the cylinder structure. In this regard, the printing of cylinders from 4% GelMA composition could become feasible when appropriate adjustments during printing are put in place.

4. **Selection of the bionk matrix**

To select the most appropriate formulation to print the SMT, a few constructs (n>3) were prototyped by using two of the most promising formulations according to printability results (see Section 3). The selected formulations were: GelMA 8% - NaAlg 7% and GelMA 4% - NaAlg 11%. The performance of these materials in realizing the complex structure as foreseen by the SMT design (Figure 1 on the main manuscript) was compared to that of a commercial bioink formulation based on nanocellulose (CELLINK-RGD, CELLINK). Since the prototyped designs included the channels, the aforementioned inks were co-printed with the sacrificial ink (Pluronic F-127), whose printing process was optimized through the printing of serpentine designs (**Figure S11**). The printing of Pluronic F-127 was optimized to reach a line width of 200 µm circa (201.3 µm ± 19.5) on the straight pathways. (**Figure S11A, B,** and **Table S2**). Printing pluronic and the bioink in the absence of cells led to the generation of a compact matrix containing channels whose size matched one of the optimized strand widths of the sacrificial ink (**Figure S11C** and **D**).

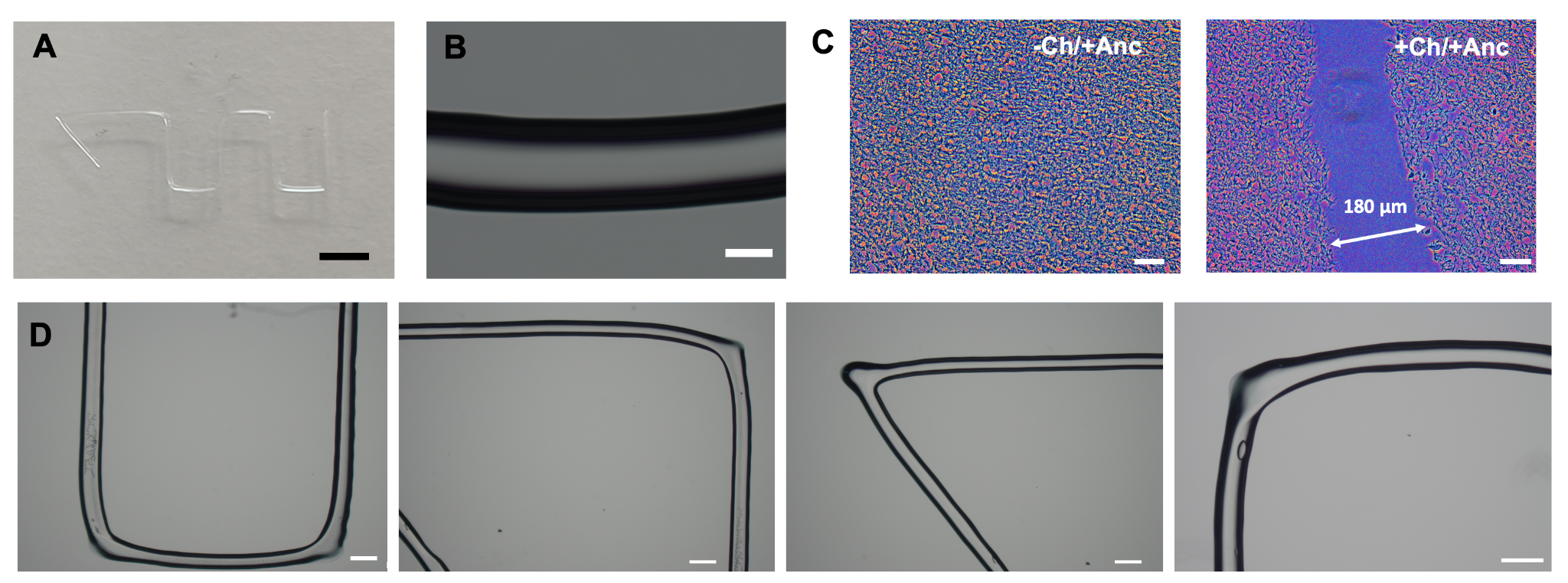

**Figure S11. Optimized printing of Pluronic F127.** Optimized printing of Pluronic: serpentine design (**A**) and line detail (**B**). Scale bars: 2 mm and 100 µm (A and B, respectively). (**C**) Histological picture of the channel in the perfusable tissue that was realized by combining the bioink with pluronic as sacrificial ink. Scale bar: 50 µm. (**D**) Representative pictures from the serpentine line. By performing the printing at the optimized values of printing speed and pressure, a line width of around 200 µm was achieved (average line width: 201.3 µm ± 19.5).

The constructs were processed for production of histological sections for morphological analysis (**Figures S12** and **S13**). The paraffinized sections (thickness: 5 µm) were stained for a standard histological coloration (H&E) with the aim to confer colored halos to the gel matrix and enable its visualization under the optical microscope under light transmission. The matrix visualization was further enhanced by applying false coloring as shown in **Figure S12A**. With this technique, it was possible to explore the architecture of the realized structures. After co-printing the inks with Pluronic and washing this away, internal cavities were generated. Cavities with extremities opened on the external space were considered as channels, and assumed to be empty volumes that could generate a functional, perfusable network. Printing the SMT structure with a highly printable material such as the CELLINK-RGD ink led to a high fraction of cavities with open extremities (around 80%) (**Figure S12B**). Also when using the GelMA 7% and NaAlg 8% a high number of functional channels (around 86%) could be generated, whereas the matrices based on GelMA 4% - Alginate 11% showed a smaller fraction of functional channels (around 64%), potentially due to the low printing accuracy and shape retention.

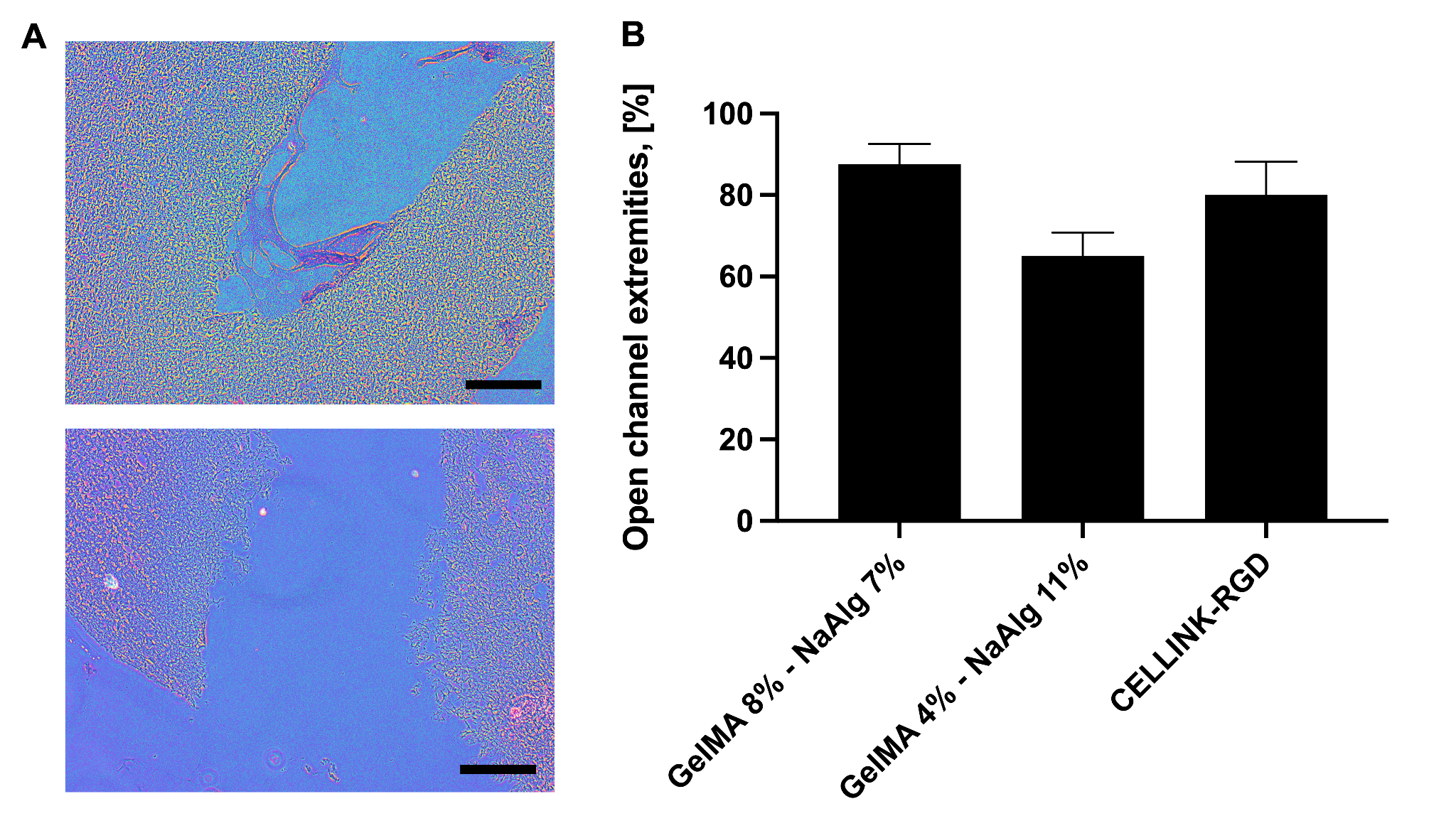
**Figure S12. Open channel extremities.** (**A**) Representative histologic section of the GelMA 8% - NaAlg 7% co-printed with alginate on the SMT construct’s design. The sporadic enclosure of the channels was observed (top) whereas the majority of channel structure was open to the external space (bottom), demonstrating structural stability. Sections were stained by Hematoxylin and Eosin staining and pictures are shown in false colors to enhance visualization of the matrix (grey) and empty areas (channel lumen, external spaces) in blue. Scale bars: 100 µm. (**B**) Percentage of channel structures that showed contiguity with the external space (open channel extremities), as quantified from image analysis on histological sections of constructs printed from two different formulations of GelMA-NaAlg and a commercial nanocellulose-based ink (CELLINK-RGD).

The morphology of the matrix realized with GelMA 8% - NaAlg 7% was further characterized via optical analysis of histological sections (**Figure S13**). Parallel cavities running along the longitudinal axis of the construct were observed (**Figure S13A**), suggesting the realization of complete channel cavities that resemble the initial design. The lumen size of such channels and the interchannel distance were measured from optical pictures at different locations within the constructs (**Figure S13B**). In particular, structural measurements were performed at different sites along the longitudinal axis of the constructs for a total explored distance of 10 mm (from -5 mm to +5 mm with respect to the median point). The lumen diameter varied in the range between 170 and 250 µm, indicating an overall homogeneous channel architecture. The negative number positions (- 5 and - 2.5 mm) corresponded to the side of the construct where the printing process started. At those levels, the constructs showed the highest diversity in channel lumen values (250 and 170 µm, respectively), whereas highly homogeneous lumen values were measured in the rest of the constructs (around 210 µm). The range of variation for the interchannel distance was contained (300-350 µm), which confirmed the realization of an accurate and regular printing scheme.

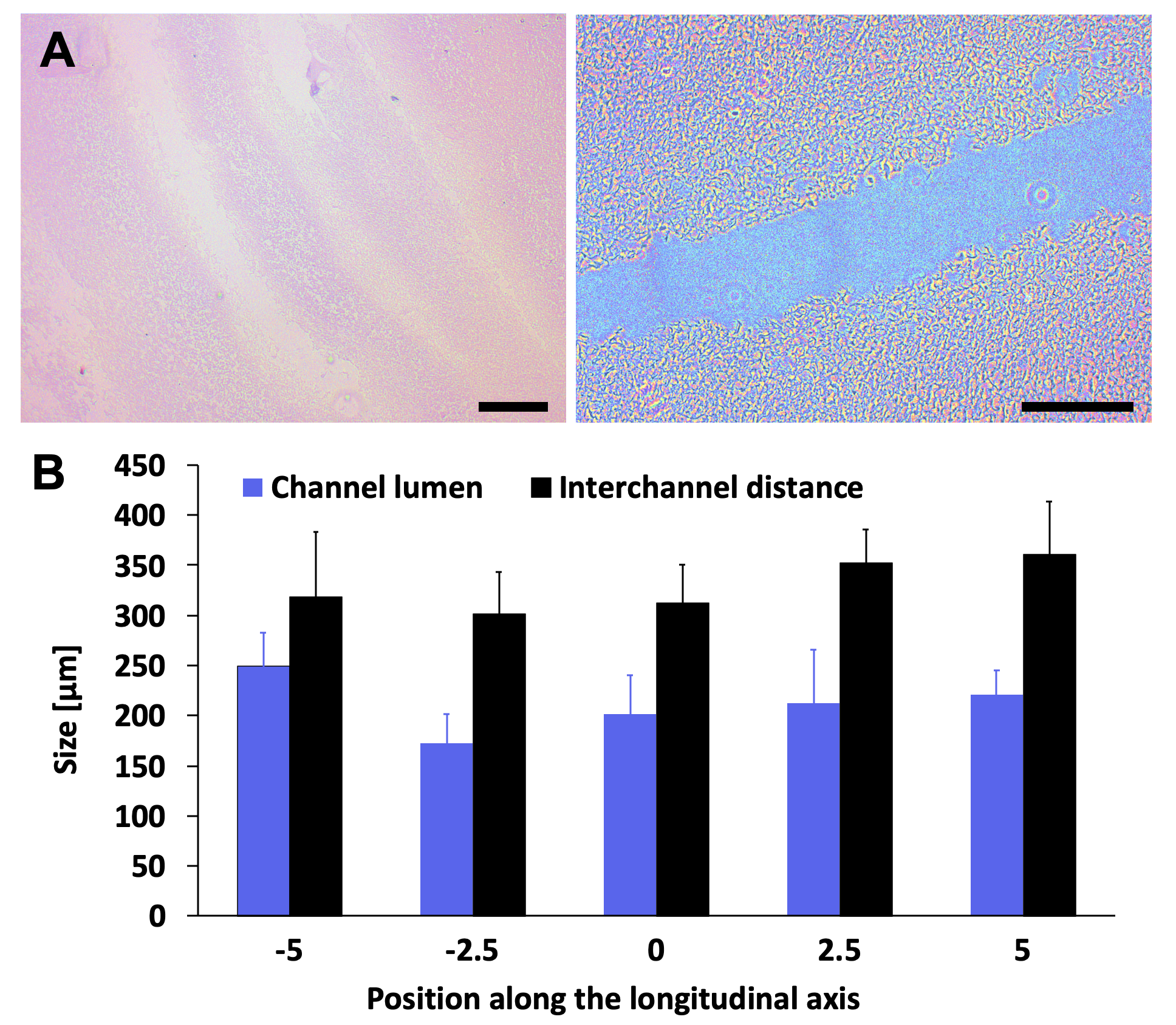

**Figure S13. Morphology of the realized channel design.** (**A**) Representative histological sections of the GelMA 8% - NaAlg 7% co-printed according to the design of the perfusable SMT. The matrix showed parallel cavities aligned with the long axis of the construct shape. Magnification of 10 and 20𝗑 (left and right, respectively). Scale bars: 300 and 200µm, respectively. (**B**) Structural characteristics of the matrix as calculated from optical analysis of the histological sections. The size of channel lumens (blue bars) and the interchannel distance (black bars) were quantified on coronal sections of the constructs at different positions over the channel longitudinal axis (from -5 to + 5 mm with respect to the median axis of the construct).

Moreover, the morphology of the scaffold was examined to determine the quality of the bioprinting outcome. The STM constructs based on GelMA 8% - NaAlg 7% displayed intact burdens as well as homogenous and compact matrices. Fractures in the polymeric matrix were only sporadically observed. On the contrary, in the constructs based on different ink formulations, fractures on the borders and in the internal areas were frequently observed (**Figure S14**).

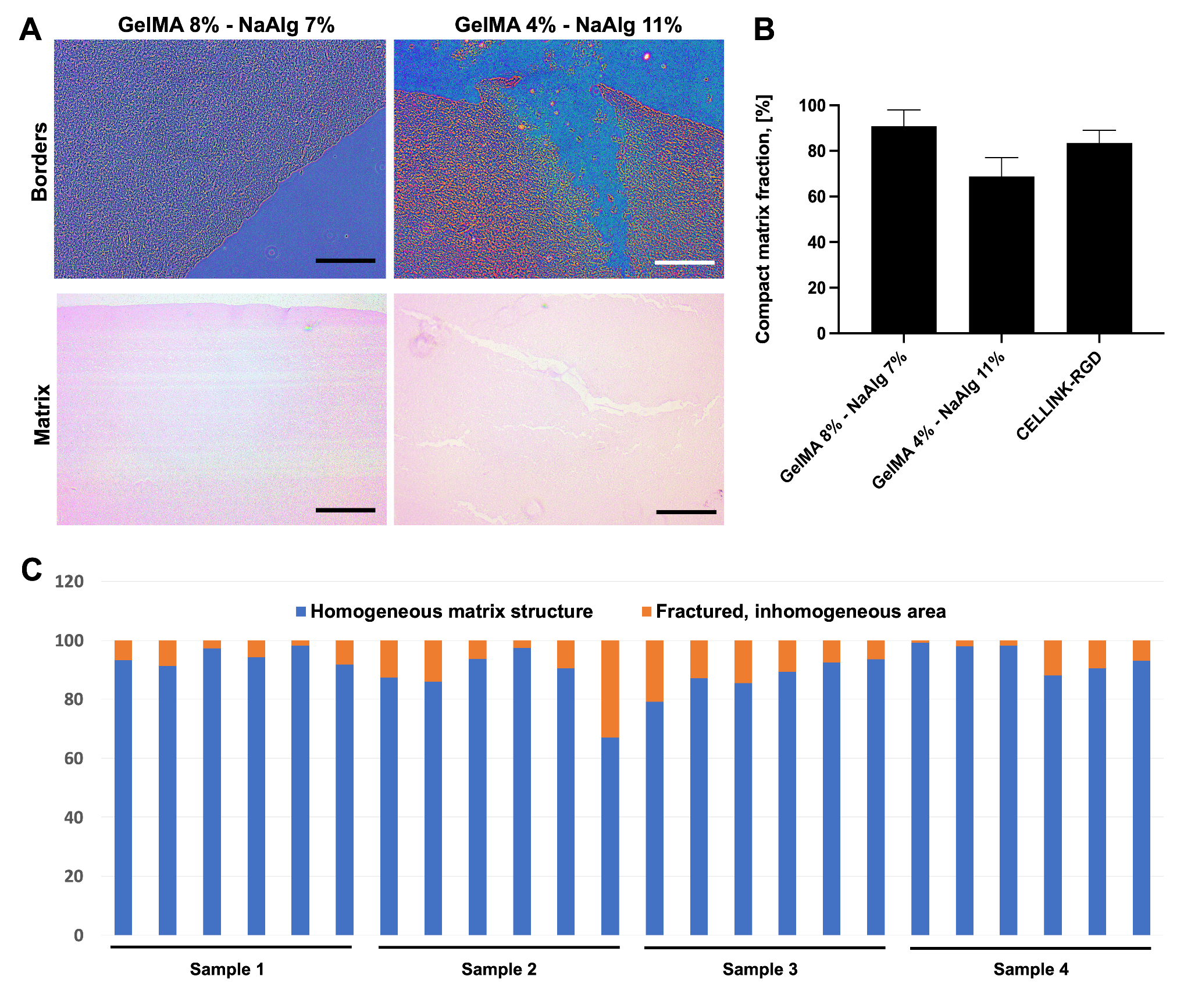

**Figure S14. Morphology of the scaffolds.** (**A**) Representative histological sections of the GelMA 8% - NaAlg 7% ink and the GelMA 4% - NaAlg 11% printed according to the design of the perfusable SMT. Borders (top) and matrix (bottom) structures are shown in false colouring and normal H&E colors, respectively. Magnifications are 20 and 10𝗑 (top and bottom, respectively); scale bars are 100 and 200 µm (top and bottom, respectively). (**B**) Percentage fraction of the homogenous, unfractured matrix as quantified by image analysis of the histological coronal sections of the SMT constructs. (**C**) Representative values of the intact and fractured matrix for four samples of SMT realized from the GelMA 8% - NaAlg 7% ink.

5. **Bioactivity on the bioink**

Cell bioactivity on the printed scaffolds was observed at different time points during culture (days 3, 6, and 15) (**Figure S15**). The scaffolds were analyzed via the Live/Dead assay on a confocal microscope and pictures were acquired at two different tissue depths below the surface (100 and 500 µm, approximatively), and on the border regions. The stainings revealed viable constructs for all the investigated conditions. Overall high cell viability was verified at deep regions in all of the constructs, with however more cells staining red (so dead cells) in the constructs lacking channels. Cell morphology (as revealed by the green staining) appeared to be elongated or myotube-like in the constructs exposed to mechanical stress, suggesting a later stage of SMT maturation.

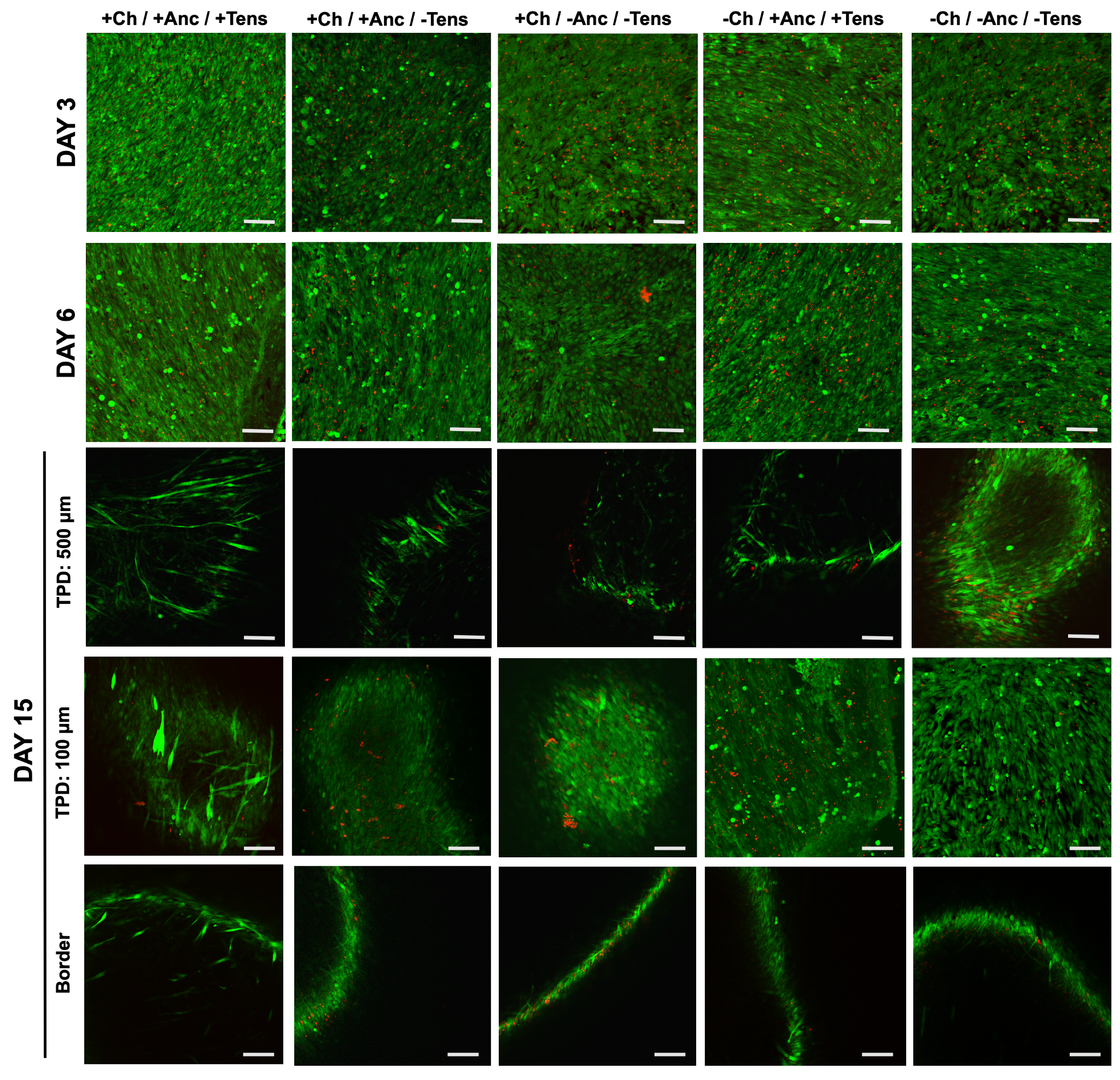

**Figure S15. Cell viability and death on maturing scaffolds.** Maturing scaffolds were stained for Live/Dead assay and pictured at different time points (3, 6, and 15 days). Matured scaffolds were pictured also at two different tissue penetration depths (TPD: 100 or 500 µm) and on the scaffold border via confocal imaging.

6. **Perfusion of the constructs**

To simulate tissue perfusion by the cell culture media, the constructs were stained repeatedly with a solution based on cell culture media added with a Red dye Erythrosin B (0.1% w/v; Sigma-Aldrich; product number: 200964). Briefly, the constructs were put on a tilted glass surface (30 °C) and three aliquots of red staining solution were repeatedly added to the upper extremity of the construct (time interval among additions: 1.5 minutes). The staining solution was left to perfuse or diffuse the tissue constructs following gravity. The stained (%) fraction of the construct was calculated from optical pictures at different time points to describe the perfusion kinetics. Figure S16 shows the lateral view of constructs stained immediately after bioprinting. Construct with channels were rapidly stained through their entire length, regardless of the presence or the absence of the anchor structures. After a few minutes, the staining solution was visible all over the tissue block surface, suggesting complete perfusion of the whole tissue amount. On the contrary, in the same time range, the non-channeled construct remained unstained, and after the last addition, the staining solution percolated around the construct down alongside the tilted surface. This observation suggested that the staining solution could only accumulate on the deposition site and could not effectively penetrate the tissue structure, even in the absence of the anchor structure.
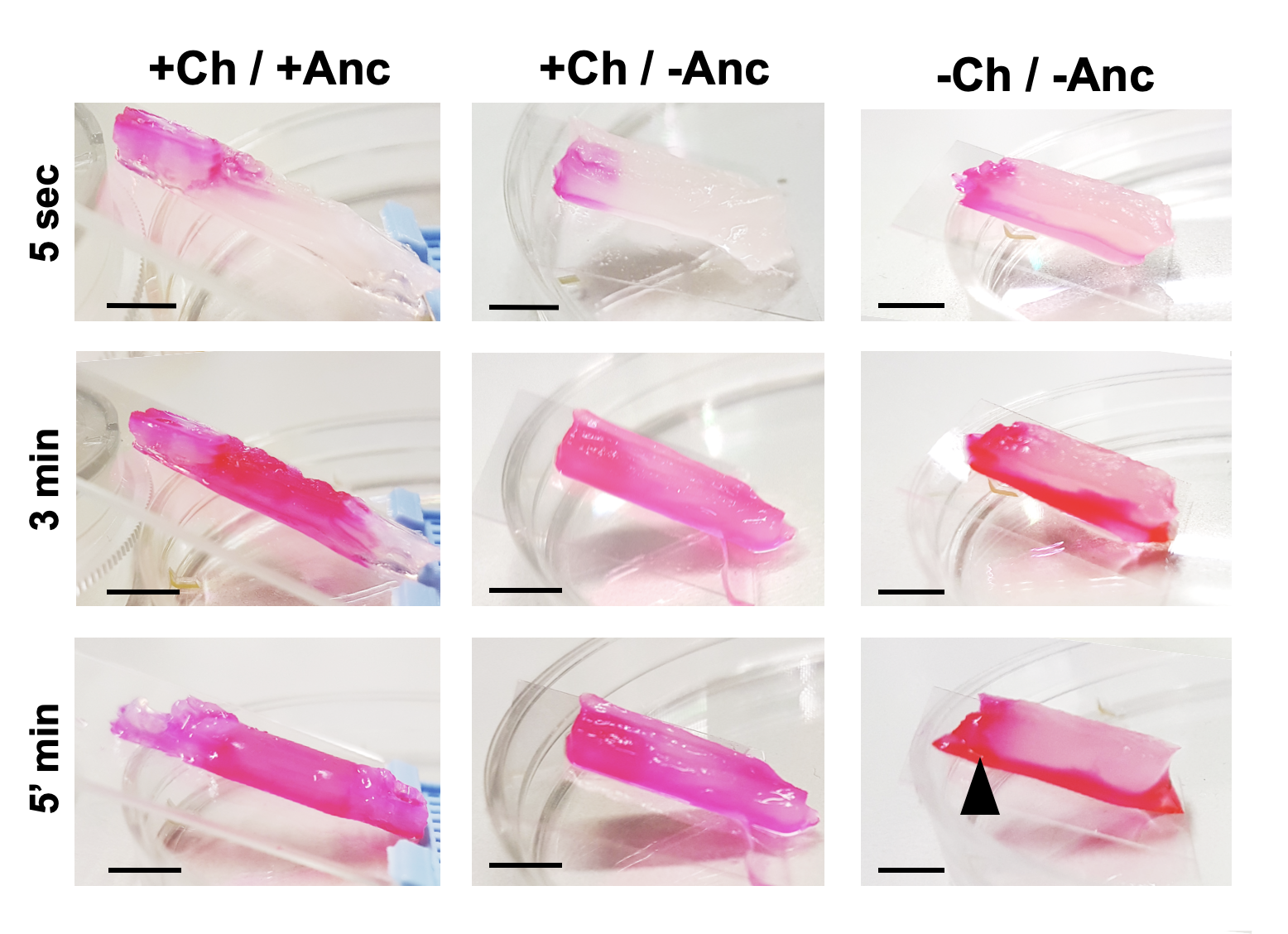

**Figure S16. Perfusion of scaffolds after bioprinting.** Representative optical pictures (lateral view) of the perfusion in the constructs immediately after bioprinting: the constructs with no channels were surrounded by the added staining solution, which could not penetrate the tissue. The solution percolated through it or slided around it (black arrow). Scale bar: 5 mm.

7. **Tissue-anchor interface characterization**

To gain insights into the structure of the realized anchors and understand how muscle tissue adapted to it, we imaged the tissue-anchor interface via two imaging techniques: confocal imaging and Scanning Electron Microscopy (SEM). Confocal imaging on day 5 revealed a dense cell population on the tissue-anchor interface area and in the matrix projections within the anchors (**Figure S17**).
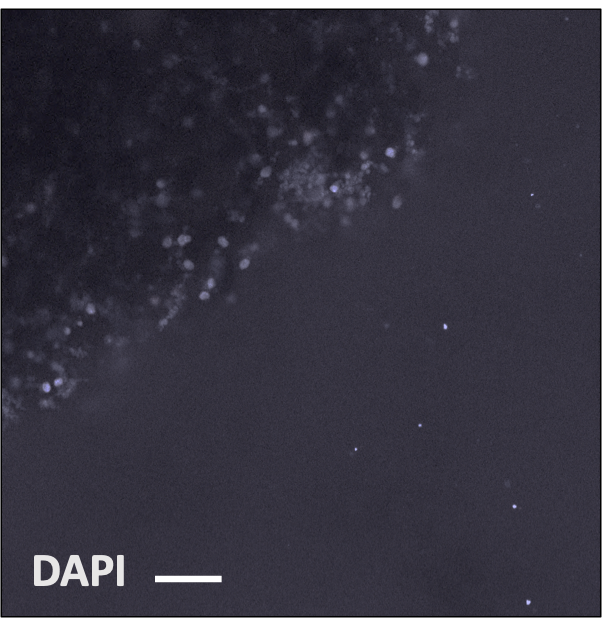

**Figure S17. Cell density at the tissue-anchor interface.** Greyscale confocal imaging of the biohybrid interface (day 5) with cells (DAPI-stained nuclei in white) growing on the synthetic surface (right side). Scale bar: 50 µm.

We observed the biohybrid interface also via SEM, focusing the analysis on the transverse section of the external anchor-tissue interface, as indicated in **Figure S18A**. Cells grew to form a biofilm that adhered to the anchor surface (**Figure S18B-F**) by tightly interacting with the scaled topography of the anchor surface (**Figure S18G**-**H**).

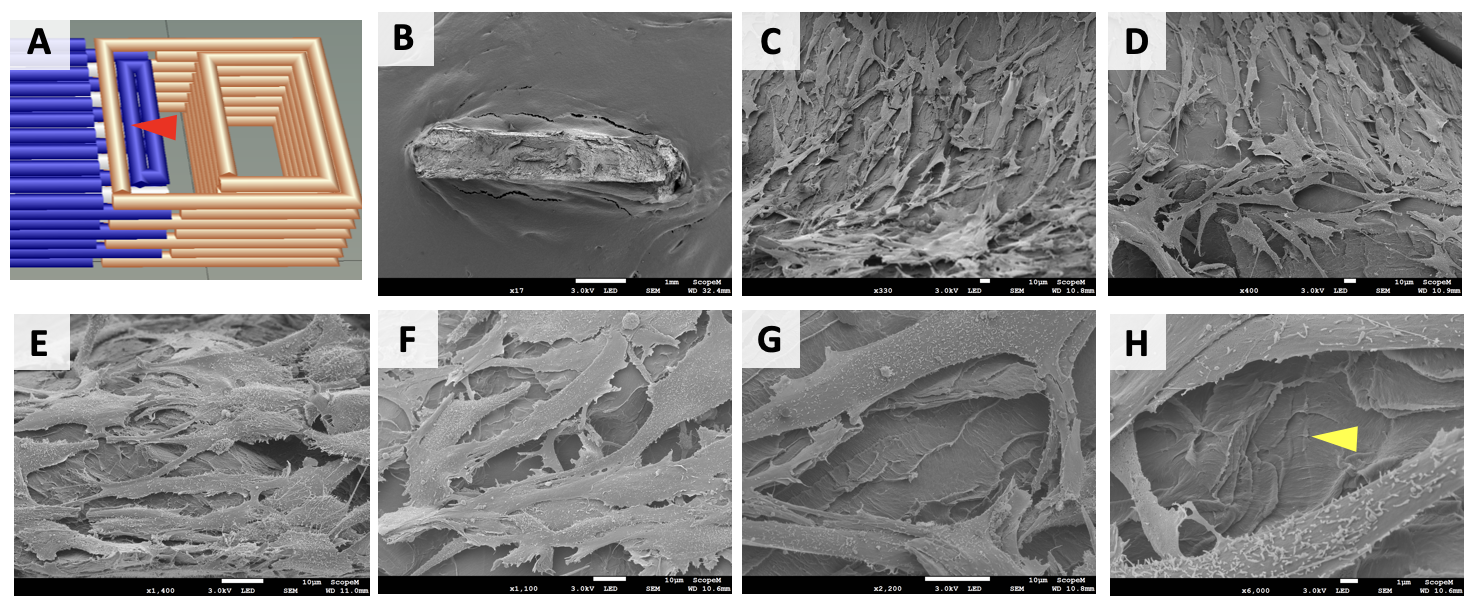

**Figure S18. Anchor-tissue external interface (central area): cell growth and adherence.** Scanning Electron Microscopy (SEM) imaging of the anchor was taken at the tissue-anchor interface within the anchor structure (red arrow in **A**) at day 15. Imaging the transverse surface of the anchor (top layer, **B**) revealed cells adhering to surfaces (**C** and **D**) to form stacked cell layers (**E-F**). Cells spread to adhere to the scaly topography of the anchor substrate (yellow arrows in **G-H**).

Cells displayed a myoblast-like shape and their membrane was widely covered with filopodia (**Figure S19**).

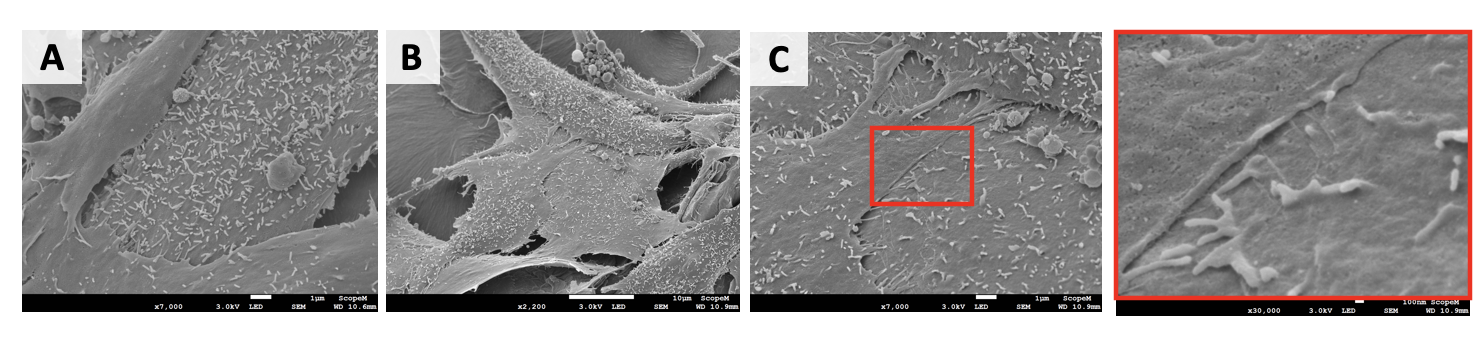

**Figure S19. Anchor-tissue external interface (central area): cell morphology.** Scanning Electron Microscopy (SEM) imaging of the cells showed their membrane morphology featuring filipodia and many newly born membrane extroflections suggesting cell bio-activity (**A-C**). Red squares indicate the inset areas showing magnified detail of myoblasts (right picture in **C**).

Dense biofilm formation was observed on the left side of the anchor transverse section, where cells followed the structural architecture of the anchor (**Figure S20**).

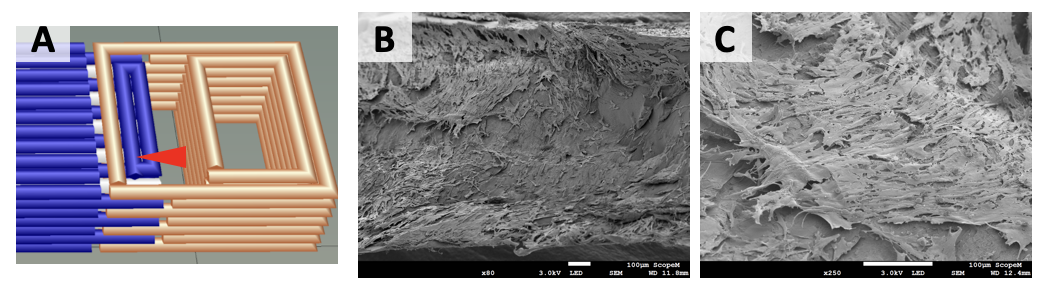

**Figure S20. Anchor-tissue external interface (left side): cell growth and adherence.** Scanning Electron Microscopy (SEM) imaging of the anchor was taken at the tissue-anchor interface within the anchor structure (red arrow in **A**) on day 15. Imaging of the transverse surface of the anchor (top layer) revealed dense muscle tissue attached to the anchor substrate (**B**) with cells adhering to the anchor surface (**C**).

In this area, the cells were compacted in a densely packed biofilm (**Figure S21A**-**B**) where cells extended their filopodia (**Figure S21C**-**D**), and were tightly adherent to each other, suggesting the occurrence of either cell proliferation (*i.e*., cellular division) or cell fusion for myotube formation (**Figure S21E**).

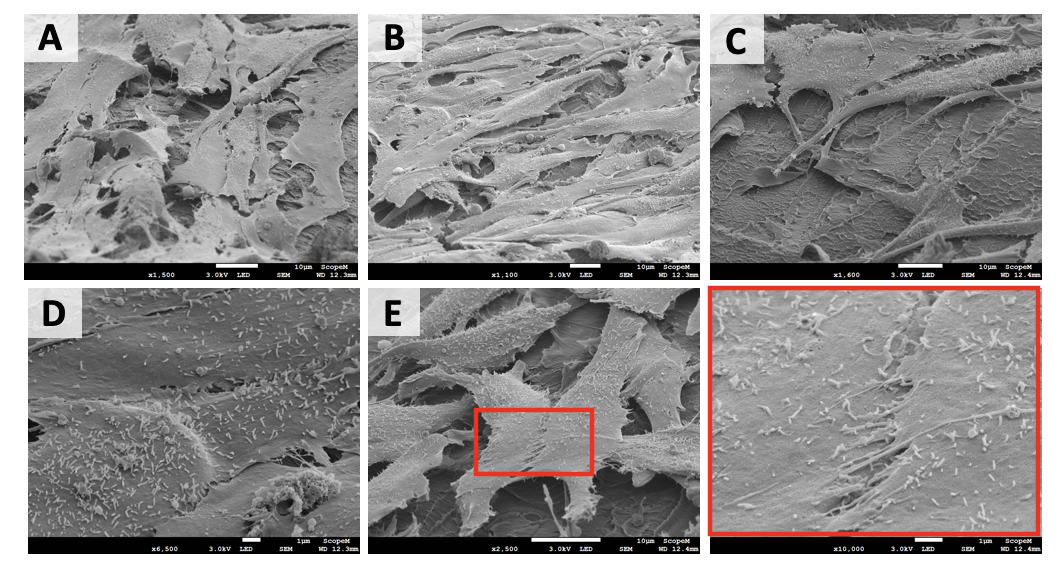

**Figure S21. Anchor-tissue external interface (left side): cell morphology.** In the muscle tissue, cells were densely compacted (**A-B**). The anchor surface was visible under the cells (**C**). Filopodia and cilia were visible on the cell membrane (**D**) and signs of tight cell-to-cell adherence were observed, suggesting potential myoblast cell fusion (**E**). Red squares indicate the inset areas showing magnified detail of the fusion interface (**E**).

Examining the bottom surface of the anchor (**Figure S22A-D**), residuals of the muscle tissue from the layer below were observed, revealing a dense myoblast population growing without any homogeneously defined directionality (**Figure S22E-H**).

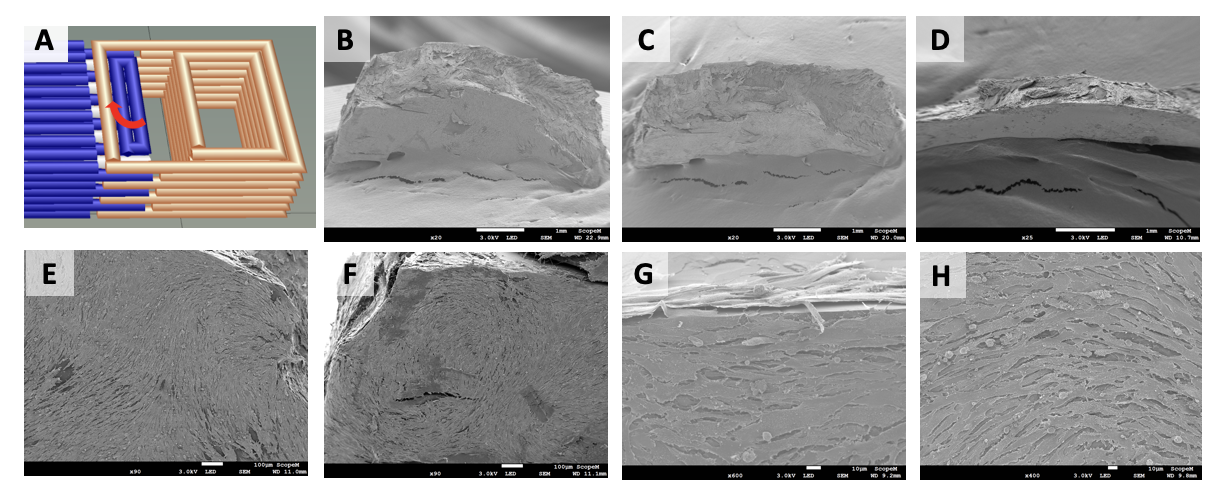

**Figure S22. Anchor-tissue external interface (second layer): muscle tissue formation.** Scanning Electron Microscopy (SEM) imaging of the anchor was taken at the tissue-anchor interface within the anchor structure (red arrow in **A**) on day 15. Imaging the bottom surface of the anchor (top layers, **B-D**) revealed muscle tissue attached to the anchor surface (**E-H**).

In the biofilm, fused cells were observed (**Figure S23A**), as well as sporadic rounded bodies (potentially apoptotic bodies, **Figure S23B**). Disrupting the tissue layer revealed the anchor surface below (**Figure S23C**).

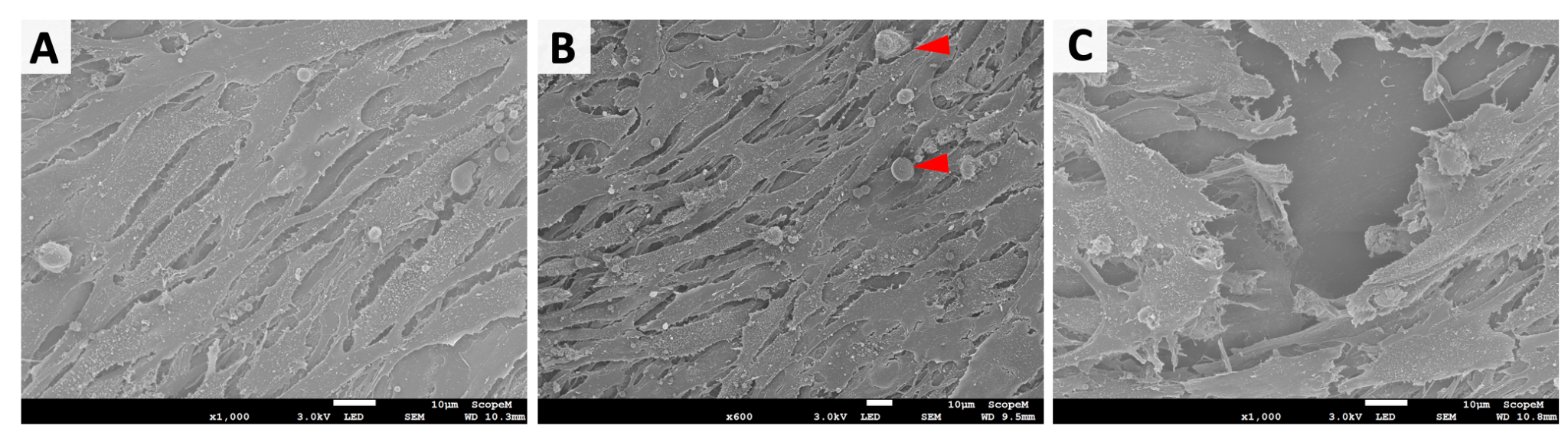

**Figure S23. Anchor-tissue external interface (second deposited layer): muscle tissue.** SEM imaging reveals the formation of a dense biofilm, in which cells proliferated and started fusion (**A**). Bodies with rounded morphology (**B**). Disrupted biofilm reveals the anchor surface beneath (**C**).

In the muscle layer, cells showed a myoblast-like morphology and a bio-active membrane that generated and extended filopodia of different sizes for cell-cell and cell-surface contact (**Figure S24**).

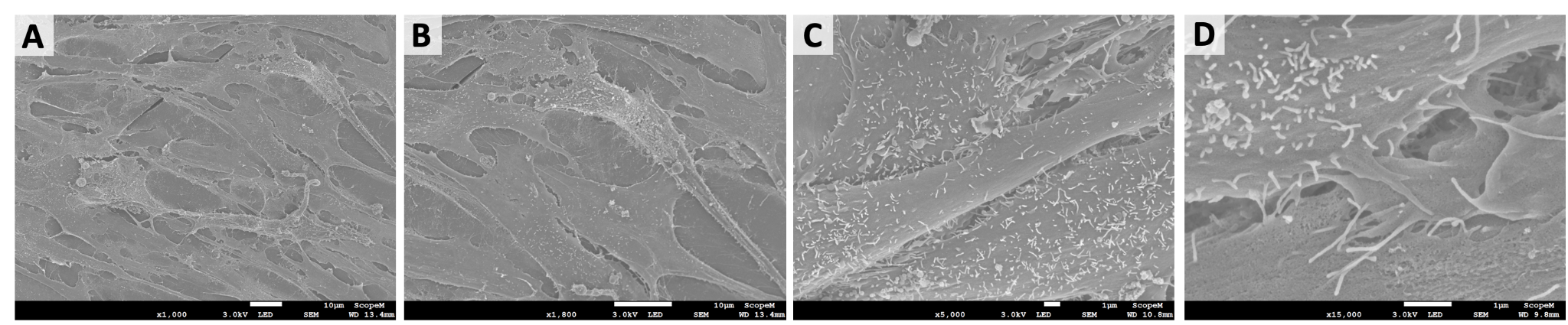

**Figure S24. Anchor-tissue outer interface (second deposited layer): cell morphology.** SEM imaging showed cells with a myoblast-typical morphology (**A** and **B**) with cell membranes covered with filopodia of different sizes (**C** and **D**) that established contact with other cells and the surrounding environment.

On the external corner side of the anchor, the cell density was low (**Figure S25**). Overall cells in this area displayed a fibroblast-like morphology suggesting the presence of adherent myoblasts on the anchor surface.

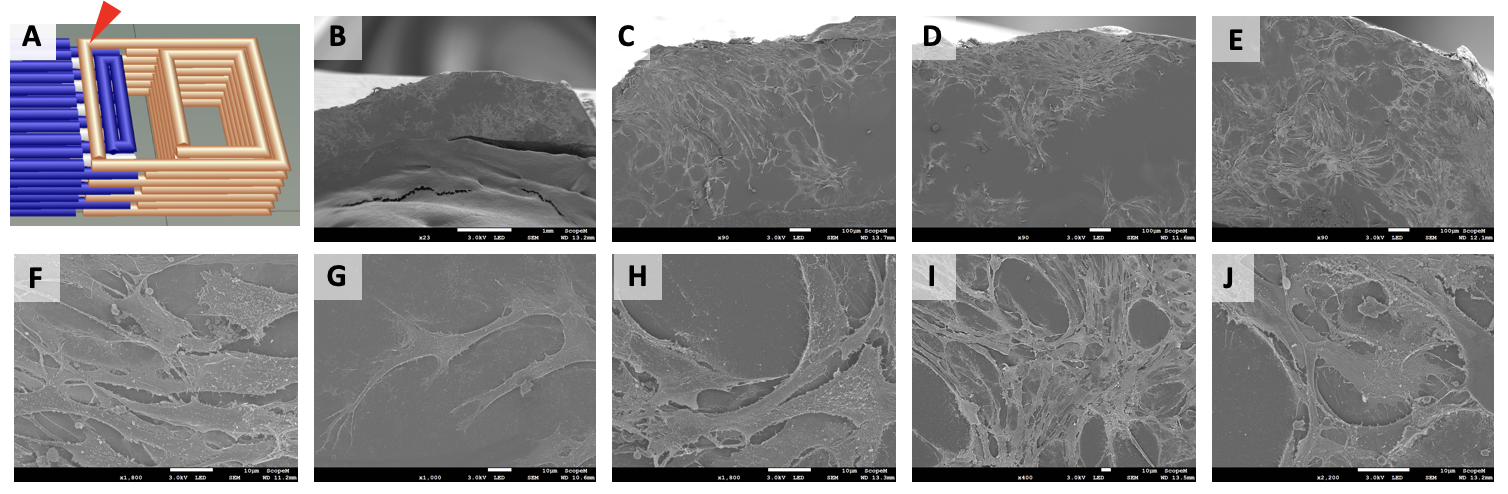

**Figure S25. Anchor-tissue outer interface (anchor corner): cell spreading.** SEM imaging of the anchor was taken on the anchor structure in the corner area (red arrow in **A**) on day 15. Imaging the external anchor surface from the side (top layer, **B-E**) revealed the spreading of cells with myoblast-like morphology that adhered to the anchor material mostly forming a non-dense monolayer (**F-J**).

In this area, cells showed a myoblast-like morphology and a bio-active membrane that generated filopodia of different sizes and extended them to establish cell-surface contact (**Figure S26**).

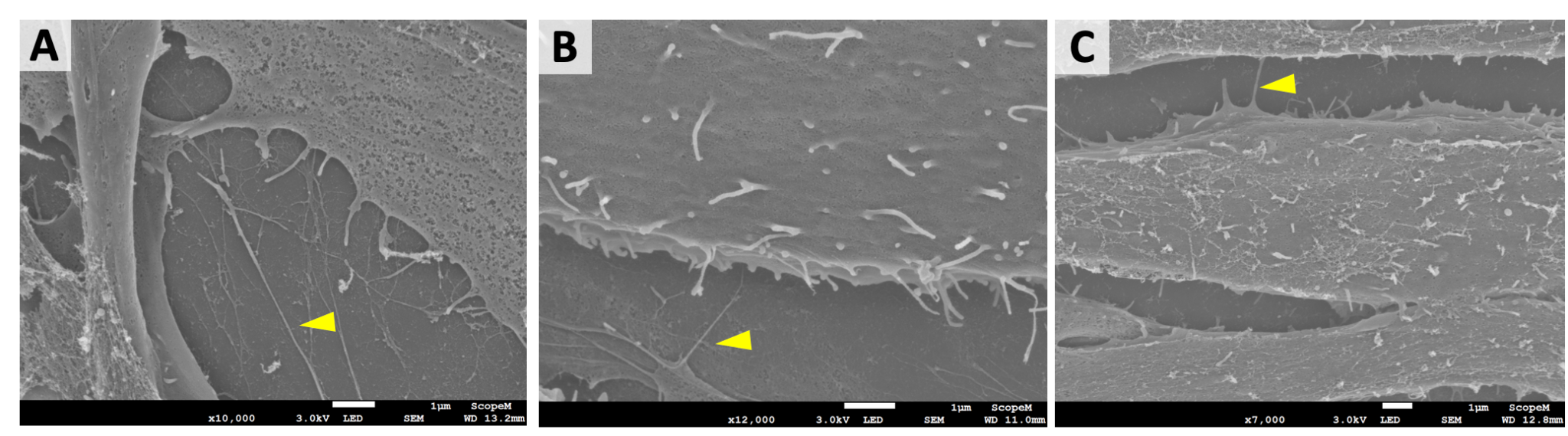
**Figure S26. Anchor-tissue outer interface (anchor corner): cell morphology.** SEM imaging of the cells reveals their adhesion sites to the anchor surface. Elongated formations indicated filopodia (yellow arrows) that sprouted from the cell membrane and mediated cell-anchor attachment.

When examining the bottom side of the later anchor part, SEM revealed muscle tissue of the second deposition layer (**Figure S27A-B**). The tissue followed the anchor architecture while preserving functional structures such as empty chambers deriving from tissue perfusable channels and their connection to the anchor structure (**Figure S27C**). In this area, cells with different morphology referring to various states of myogenesis were observed, including spread myoblasts (**Figure S26D**), spindle-shaped myoblasts (**Figure S26E**), and fused myoblasts (**Figure S26F**).

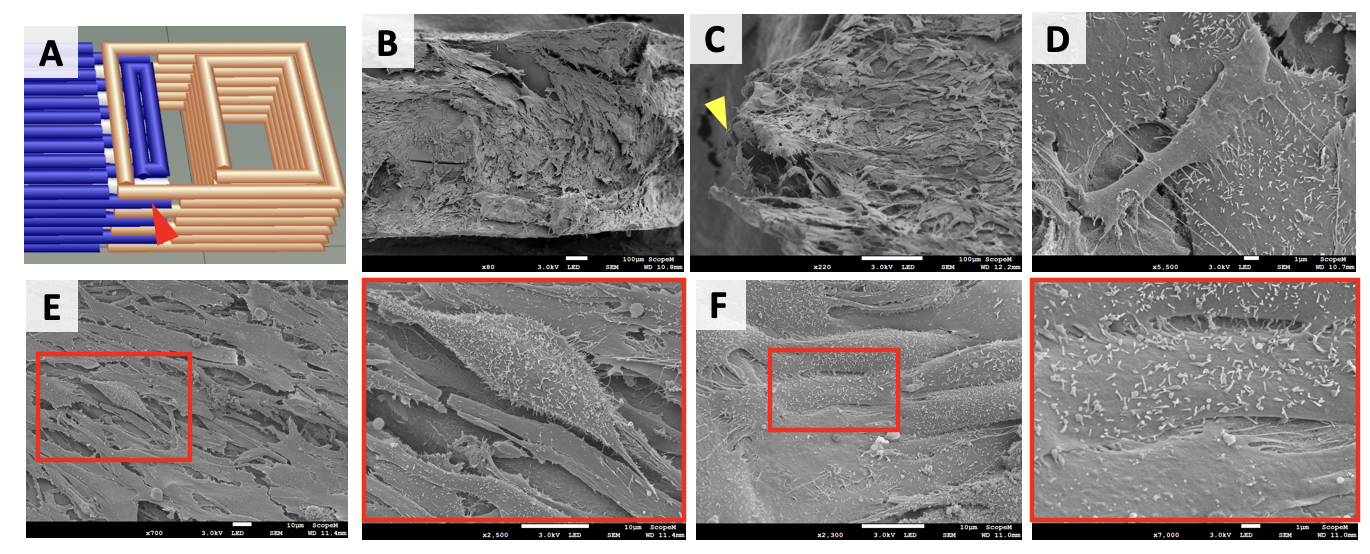

**Figure S27. Anchor interlayer space: muscle tissue.** SEM imaging was taken in the interlayer space of the anchor structure (red arrow in **A**) showing muscle tissue with a dense cell population (**B**). The tissue followed the anchor topography preserving functional architecture (*e.g*., channel connection: yellow arrow) (**C**). Cells with adherent myoblast-like morphology and evident filopodia were detected (**D**). In some tissue areas, spindle-shaped myoblasts were seen (**E**), while in other areas, cells fused together, suggesting the starting of myotube formation (**F**). Red squares indicate the inset areas showing magnified detail of myoblasts (right picture in **E** and **F**).

Cells displayed filopodia (**Figure S28A-B**). Moreover, on the anchor surface, filamentous materials were observed, which could potentially be either polymeric residuals from the bioink blend or the cell-produced extracellular matrix (**Figure S28C-D**).

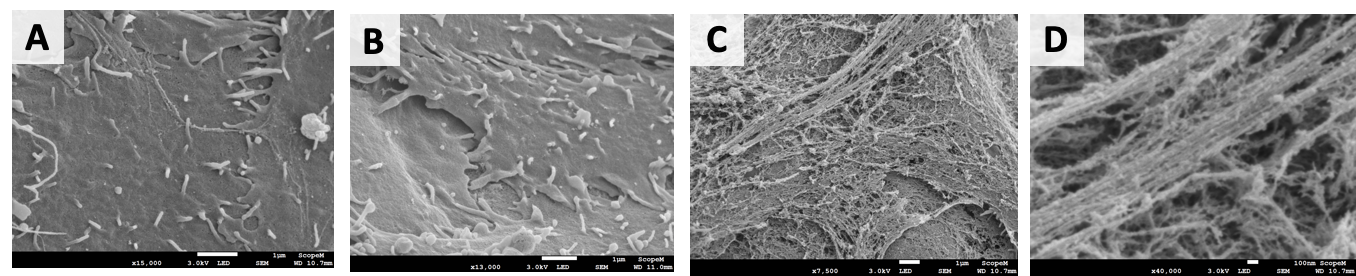

**Figure S28. Anchor interlayer space: cell morphology and matrix deposition.** Detailed SEM imaging of the cell surface revealed the sprouting of filopodia (**A-B**) and the deposition of extracellular matrix (**C-D**).

Finally, the formation of rounded bodies on the cell surface was also sporadically observed (**Figure S29**), suggesting vesicle or apoptotic bodies release.

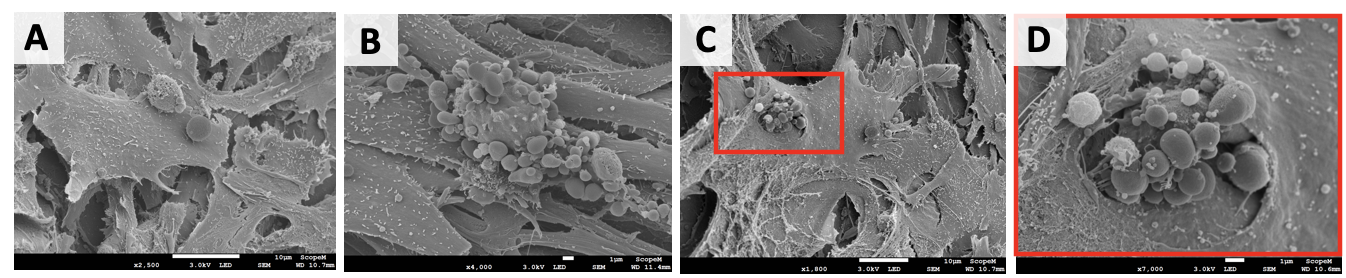

**Figure S29. Anchor interlayer space: vesicle-like bodies.** In the muscle tissue, vesicle formation on the cell surface was sporadically observed, suggesting the potential formation of apoptotic bodies (**A-C**). Red squares indicate the inset areas showing magnified detail of vesicular productions (**D**).

8. **Structural stability of the constructs**

To confer mechanical tension for muscle maturation, the constructs were fixed during the differentiation phase to agar beds at the bottom of culture wells.^6,7^ The overall stability of the biohybrid constructs was assessed by quantifying those constructs that remained intact over all the culture time by macroscopic observation (**Figure S30**). The constructs that were considered non-stable were classified into two major categories. The first category included those constructs that were seriously compromised, in which the SMT, the anchor structures or their interface were destroyed or damaged so that mounting on the maturation template was not possible. The second category reported on minimally damaged constructs, in which even though macroscopic defects were visible at the level of the SMT or the anchors, these damaged areas did not alter the assembly of the different components, and anchorage to the pillars of the maturation template was feasible. In the minimally damaged constructs, the tissue maturation process was expected to take place majorly.

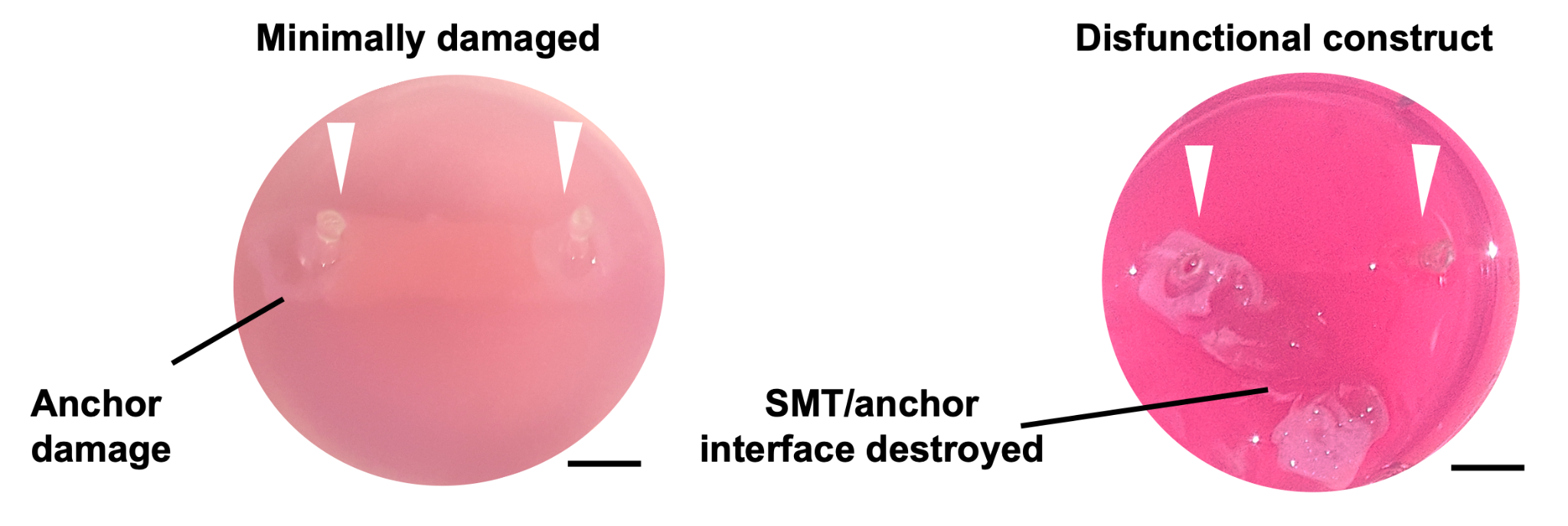

**Figure S30. Scaffold integrity evaluation.** Representative optical pictures of the GelMA 8% - NaAlg 7% constructs during culture. A minimally damaged construct (left) displays dissection of the anchor structure, but remained stably anchored on the maturation template. Ac completely destroyed biohybrid construct (right) disassembled during culture, detaching from the maturation stage. White arrows indicate the pillars’ position. Scale bars: 5 mm.

The passive force exerted by the muscle constructs on the pillars during tissue development was estimated via a custom-made cantilever system. The culture template composed of an 8mm-tick layer of agar gel (1.5% w/v agarose, Sigma-Aldrich) and filled-in P10 pipette tips was built in a petri dish (**Figure S31**). The dish was then fixed in a vertical position to orient the pillars horizontally. Pressure was applied to the central part of the tip to achieve a later displacement of the pillar of about 1.5 mm (which was observed on real constructs). The whole set-up was placed on a scale, and weight values were taken before and after the application of pressure. The force value was estimated from the weight difference between the two conditions.

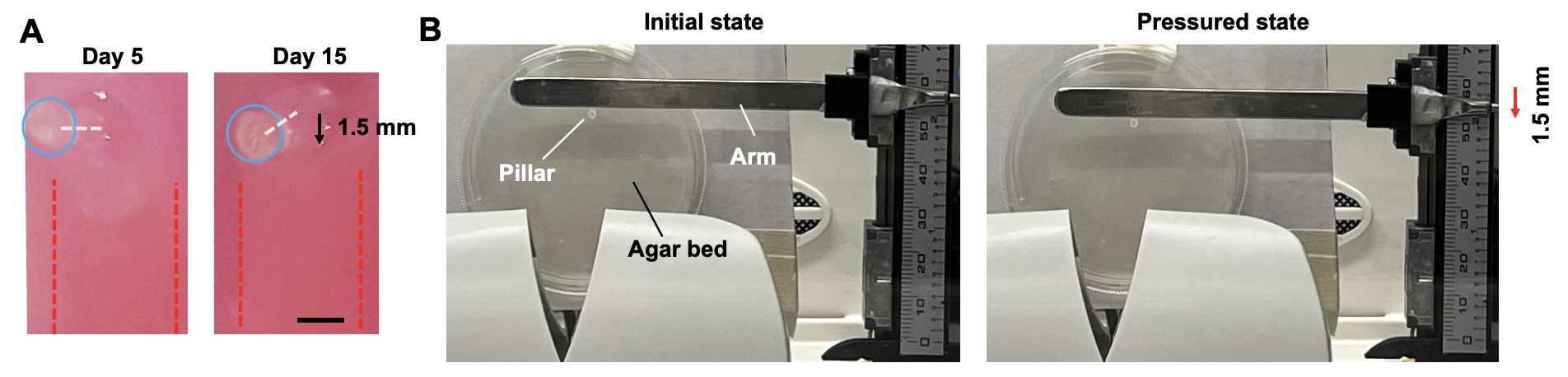

**Figure S31. Setup for passive force estimation.** (**A**) Optical pictures of anchored constructs at day 5 (first day on the maturation template) and day 15 (end time point of culture). Red dashed lines indicate the borders of the SMT; blue circles indicate the top part of the pillars; white dashed lines indicate the long axis of the pillar. Scale bar: 3 mm. (**B**) Optical pictures of the set-up used to estimate the force acting on pillars during the maturation of the biohybrid SMT construct. A layer of agarose was placed in a vertical position, and the pipette tips (acting as pillars) were inserted in the substrate and sat in a horizontal position. A rigid metallic arm was used to apply the pressure in a controlled manner, by matching specific values of lateral displacement (1.5 mm).

9. **Final remarks**

*9.1 Contribution to the design of biohybrid SMT.*

Biohybrid fabrication strategies are needed to approach the engineering of larger and more biomimetic SMT constructs that could serve in various applications, such as the treatment of large muscle defects and performant bio-actuators for robotics.^8,9^ However, the inherent constraints of biofabrication limit the scalability of SMT size beyond the cm-range, which in turn limits its actuation performance.^8,10^ SMT biofabrication should enable perfusable structures with geometries that comply with the structure of the native tissue, such as microchannels aligned in parallel to the myofiber orientation. When engineering SMT another crucial aspect to consider is tissue anisotropy, as the alignment of the muscle cells and the uniaxial myotube formation are essential for appropriate myofiber maturation, oriented contraction, and force production.^8,11,12^ Biofabrication that aligns muscle cells to form uniaxial myotubes is essential for myofiber maturation, oriented contraction, and force production.^8,11,12^ The maturation of SMT engineered at the mesoscale (*i.e*., cm size ranges) requires its development under mechanical stress, and for this purpose, the living tissue can be combined with synthetic culture templates that generate passive tension in response to the matrix shrinkage.^9,13–15^ In addition to enabling the mechanical stimulation of cells within thick constructs, biohybrid designs are needed to realize biohybrid robots, in which the SMT integrates with synthetic elements that convert its contraction into specific types of motion or locomotion.^14–18^ The design of the synthetic elements also contributes to realize the desired motion patterns.^14–18^ Anchoring synthetic structures at the extremities of thick muscle tissue blocks generates modular designs that facilitate the handling of fragile hydrogel-based 3D SMT constructs.^19^ Biohybrid fabrication can also model the myotendinous axis, for which the crucial aspect is to realize seamless interfaces to optimize force transfer from muscle to tendon.^20^ For example, Ladd et al. realized a myotendinous junction model by culturing the cells (myoblasts and fibroblasts) on a dual scaffold featuring electrospun polymer gradients that aim to replicate the regional differences of mechanical stiffness found at the interface between the tendon and skeletal muscle.^21^

In our work, we showed that layer-by-layer multimaterial bioprinting of living and non-living materials can realize large, perfusable biohybrid SMT designs that enhance tissue maturation *in vitro*. Through multimaterial bioprinting, we integrated structures with different mechanical properties into one. The living tissue was crossed by a network of perfusable microchannels oriented parallel to the main axis for fiber formation. The channel network remained contiguous with the external environment even in the presence of the anchoring structures. Moreover, the channels structure and interchannel distance remained regular over time, suggesting that the morphological changes naturally happening during tissue maturation did not compromise the functionality of the perfusable architecture.

In our system, synthetic materials and biomaterials were shaped into one during the bioprinting process. The synthetic structures were stabilized via crosslinking at the same time as the biomatrix itself. Then, the muscle tissue matured while being in direct contact with their surfaces, allowing the soft tissue to adapt and tightly adhere to the complex morphology of the anchors. Fabricating biohybrid structures in one process is likely to support the formation of coherent interfaces, which perspectively would help for improved performance during the contraction of the generated actuator.^13^ Future studies should address if our one-go fabrication process is advantageous not only to support the SMT development but also their dynamic functions, for example by increasing the efficiency of force transmission through the tendon-like structures.

*9.2 Contribution to the materials for SMT fabrication*

Thus far, only limited research focused on selecting materials and biofabrication approaches for an efficient biohybridization of engineered skeletal muscles.^22,23^ These materials should display mechanical properties that guarantee muscle cell differentiation and proper SMT formation. Much research has been done to elucidate the role for matrix or tissue stiffness in striated muscle development,^24,25^ finding that muscle cells have complex responses to the mechanical properties of scaffolds. For example, myogenic differentiation in culture depends intimately on optimal outside-in signaling mediated by the matrix elasticity.^26^ Cell fusion into myotubes might occur independent of substrate flexibility, but myosin/actin striations only emerge in scaffolds displaying stiffness that match the values of actual muscle (passive Young's modulus of~10-12 kPa).^27–30^ Unlike sarcomere formation, the cell adhesion strength augments monotonically versus substrate stiffness with strongest adhesion on stiff scaffolds.^26^ So for successful development of SMTs requires scaffold materials that display a right balance between elasticity and stiffness.

In our work, we developed a printable bioink based on GelMA and NaAlg. GelMA is a gelatin derivative hydrogel with chemical UV-based crosslinkage and excellent thermal stability. GelMA features all biofabrication-related advantages of gelatin, such as high biocompatibility, solubility in water, biodegradability, and low antigenicity. GelMA retains the native arginine-glycine-aspartic acid (RGD) peptide sequence, as well as the matrix metalloproteinase (MMP) degradation sequence, thus cells can adhere, proliferate, differentiate, and also remodel the surrounding matrix by enzymatic digestion. We added NaAlg to the bioink to modulate the rheological properties for a suitable stiffness and viscosity needed in bioprinting.

We developed our bioink formulation based on GelMA and NaAlg from an optimization study that took into account various formulations with variable fractions of the components. The different formulations were rheologically characterized in both crosslinked and non-crosslinked states, and their storage and loss moduli (G' and G’’, respectively) were determined over a wide shear stress range (**Figure S2** and **S3**). All the crosslinked samples exhibited a shear-thinning behavior, evidenced by decreasing viscosity at increasing shear rates. In fact, the G' values fell below the increasing G'', suggesting that all blends acquired a more liquid-like behavior at higher shear rates (**Figure S3**). Such a shear-thinning behavior is desirable, as it enables the material extrusion at low-pressure ranges (between 40 and 80 kPa), which do not hamper cell viability. In the formulations containing 6% GelMA or 7% NaAlg, augmenting the NaAlg or GelMA resulted in a coherent increase of the complex viscosity (**Figure S4**). In addition to exhibiting a shear thinning behavior, all the tested formulations were printable (**Table S2**). The 8% GelMA - 7% NaAlg formulation displayed the highest printability as shown on the strand assessment, accuracy, and shape fidelity tests (**Figure S4-S10**).

During the optimization, it was noticed that the total polymer concentration of the bioink blend was the primary determinant for printing accuracy. The formulations with overall polymer concentrations approximating 15% w/v displayed a high printing accuracy (>90 %), whereas 13 and 11% w/v polymeric concentrations caused the printing accuracy to decrease to approximately 60 and 50%, respectively. The lowest accuracy values were observed in the formulations containing lowest GelMA amounts (4% w/v), suggesting that GelMA is more critical for printing accuracy and shape maintenance than NaAlg. With a printing accuracy of above 90%, the 8% GelMA - 7% NaAlg ink formulation was selected to prototype the channeled constructs first without and then with cells.

With this ink, we could realize centimeter-scale perfusable SMT constructs consisting of a multilayered tissue structure with internal microchannels and stably interfaced with external anchoring structures. As seen in axial and coronal histological sections of the cell-laden constructs after bioprinting, channels were present in the SMTs that were co-printed with Pluronic, whereas the matrix structure of controls with no channels was homogenous, regardless of co-printing with anchoring structures. We demonstrated efficient perfusion of the intra-tissue channels in the presence of the anchoring structures that were designed to allow for continuous flow between the channel lumen and the external environment. The channels enabled efficient medium exchange through all the tissue and was found in good correlation with the indexes of cell viability. In addition, as the channels were biomimetically designed to run with a parallel orientation to the longitudinal myofiber axis, their presence played also a role in contact-guided cell alignment and the unidirectional growth of the fibers. As seen in axial and coronal histological sections of the cell-laden constructs after bioprinting showed channels in the SMT that were co-printed with Pluronic, whereas the matrix structure of controls with no channels was homogenous, regardless of co-printing with anchoring structures.

With this work, we demonstrated the efficient realization of SMT through a bioink based on GelMA and NaAlg, in the absence of the materials that are typically used for SMT maturation. These materials are often expensive or not suitable for bioprinting. As an example, Matrigel is a highly expensive material that requires specific temperature conditions for printing, which limit its combination with other materials useful for biohybrid designs. Note that the differentiation of C2C12 cells and the formation of functional fibers in common biomaterials is achieved within one week.^15,61–65^ In our system, we could achieve SMT maturation only exposing the cells to more days of differentiation stimuli (11 days), and this followed an initial phase required for cell proliferation (4 days).

*9.3 Understanding the muscle tissue maturation*

Each step of the myogenic differentiation is regulated by muscle-derived cytokines, and several cytokines contribute to more than one stage of this process. To understand tissue maturation in our system, we quantified the production of cytokines, which are well-known autocrine regulators of muscle differentiation. Even if the working mechanism of their myogenic function is not always well-defined, several cytokines have been proved to exert an autocrine regulation on developing skeletal muscle.^31^ To better understand tissue maturation in our system, we quantified the production of cytokines that are well-known autocrine regulators of muscle differentiation. Bio-factors supporting tissue maturation were exponentially expressed for the duration of culture. Mechanical tensioning and switching from growth to differentiation media lead to a decrease of BMP-4 and Myostatin in the second half of culture time, which fostered myogenic differentiation. The absence of channels affected the production of severals cytokines (such as BDNF, VEGF, IL-1, IL-6, and IL-12) to a different extent depending on the presence or absence of the passive tension. Co-printing perfusable SMT with anchors was found not to have evident effects on the cytokine production, leading to no secretome differences between the “+Ch/+Anc” and “+Ch/-Anc” conditions. This observation reinforces the hypothesis that our biohybrid design preserved cell viability by effectively combining the channels of the soft tissue with the void networks of the anchors.

*9.4 Impact and future research*

Biohybrid fabrication of large, SMT constructs could improve the treatment of muscular defects and the creation of 3D cell culture models and bio-actuators for robotics.^1^ Bioengineering muscle tissue *in vitro* had so far been strongly hindered by poor shape fidelity, simple structural hierarchy, and lacking biomimicry; missing a suitable combination of biomaterials and biofabrication processes had led to adversarial structural phenomena occuring during tissue maturation. Our work is a proof of evidence that extrusion-based bioprinting can control the deposition of multiple materials in highly complex designs enabling better muscle tissue generation *in vitro*. Our bioink formulation has the appropriate mechanical properties and sufficient biocompatibility to endure an intricate fabrication and SMT maturation process. Our biohybrid SMT design has fine perfusable channels that pass through both the cell construct and its adjacent anchors to allow for an exchange of liquid media with the external environment. This design mimics natural blood vessels in their size and longitudinal orientation. With this result, we have demonstrated that layer-by-layer multimaterial bioprinting can realize large, perfusable biohybrid SMT designs that enhance tissue maturation *in vitro*. In the next few years, biohybrid bioprinting will drive the development of novel strategies in muscle tissue engineering for new applications in fundamental biomedical research, implantology-oriented bioengineering, nutrition, and biohybrid robotics.

10. **References**

31.	Waldemer-Streyer, R. J., Kim, D. & Chen, J. Muscle cell-derived cytokines in skeletal muscle regeneration. *FEBS J.* **n/a**,.
